# Endocrine therapy-specific lineage and partial epithelial-mesenchymal reprogramming defines divergent resistant cell-states in ER+ breast cancer

**DOI:** 10.64898/2026.05.11.724242

**Authors:** Sarthak Sahoo, Sejal Khanna, Swayamshree Senapati, Hemant Kumar, Jyothi S. Prabhu, Dimple Notani, Sridhar Hannenhalli, Mohit Kumar Jolly

**Affiliations:** Department of Bioengineering, Indian Institute of Science, Bangalore, Karnataka, India; Department of Biology, Indian Institute of Science Education and Research Tirupati, Tirupati, Andhra Pradesh, India; School of Basic Sciences, Indian Institute of Technology, Bhubaneswar, Odisha, India; Division of Molecular Medicine, St. John’s Research Institute, St. John’s Medical College, Bangalore, Karnataka, India; National Center for Biological Sciences, Tata Institute of Fundamental Research, Bengaluru, Karnataka, India; Cancer Data Science Laboratory, Center for Cancer Research, National Cancer Institute, National Institutes of Health, Bethesda, MD, USA

**Keywords:** endocrine therapy, therapy resistance, partial EMT, lineage plasticity, ER+ breast cancer

## Abstract

Acquired resistance to endocrine therapy remains a primary obstacle in the clinical management of estrogen receptor-positive (ER+) breast cancer. While resistance is frequently accompanied by transcriptional rewiring and lineage plasticity, how specific pharmacological modalities dictate divergent resistance trajectories remains poorly understood. Here, we integrate multi-omic profiling, spanning bulk and single-cell transcriptome, chromatin architecture (Hi-C), and the cistrome, to systematically compare the mechanisms involved in adaptive resistance to selective estrogen receptor modulators (SERMs, e.g., tamoxifen) and degraders (SERDs, e.g., fulvestrant), and the mechanism driven by constitutive ESR1 mutation, to characterize how mode of ERα perturbation influences lineage identity and epithelial–mesenchymal state. We found that tamoxifen resistant (TamR) cells occupy a distinct transcriptional state characterized by coordinated luminal erosion, partial basal lineage activation, and stabilization of a partial epithelial–mesenchymal (pEMT) program. In contrast, fulvestrant resistant (FulR) cells primarily suppress ER signaling without extensive lineage reprogramming. Finally, ESR1 mutant cells recapitulate ligand-driven ER hyperactivation with limited engagement of mesenchymal and basal gene expression programs. Chromatin profiling further revealed that SERM resistance is accompanied by higher-order genome reorganization, including A-to-B compartment switching at luminal regulators such as GATA3 and ESR1, redistribution of ERα and FOXA1 binding, and consequent activation of a pEMT program. Furthermore, we show that SERM-induced reprogramming is accompanied by a distinct mode of immune evasion where the reprogrammed cells do not engage classical T-cell exhaustion programs but instead exhibit coordinated loss of major histocompatibility complex (MHC) class I antigen presentation and establishment of a pro-tumorigenic signaling that strongly predicts adverse survival outcomes in patient cohorts. Together, these findings indicate that endocrine resistance does not converge on a single molecular endpoint but instead reflects drug-specific adaptive states defined by ER signaling context, lineage identity, and chromatin architecture. Our study establishes the basal–pEMT axis as a coordinated, epigenetically encoded module of SERM-induced plasticity and reframes endocrine resistance as a multidimensional evolutionary process shaped by therapeutic mechanisms of action.

## Introduction

Adjuvant endocrine therapy (ET) is the cornerstone of clinical management for most patients diagnosed with Estrogen Receptor-positive (ER+) breast cancer (1). Standard protocols prescribe a five-year therapeutic regimen, which may be extended to ten years based on risk stratification variables including tumor burden, histological grade, and nodal involvement. Whether utilized in the adjuvant setting to mitigate recurrence or the neoadjuvant setting to facilitate surgical resection via tumor debulking, ET relies on the precise targeting of ER signaling to arrest neoplastic proliferation. Therapeutic selection is strictly stratified by menopausal status and patient risk profiles (2). The established pharmacological armamentarium includes ovarian suppression (via GnRH agonists or surgical oophorectomy), Aromatase Inhibitors (AIs), Selective Estrogen Receptor Modulators (SERMs), and Selective Estrogen Receptor Degraders (SERDs) (3). Mechanistically, SERMs (e.g., tamoxifen) function as competitive antagonists at the receptor interface, whereas SERDs (e.g., fulvestrant) induce proteolytic degradation of the receptor to abrogate signaling. On the other hand, Aromatase Inhibitors (e.g., letrozole, anastrozole, and exemestane) target the biosynthetic pathway rather than the receptor itself. By inhibiting the cytochrome P450 enzyme aromatase, they block the peripheral aromatization of androgens into estrogens (4). This systemic blockade profoundly depletes circulating estrogen levels, thereby depriving the estrogen receptor-positive tumor of its primary activating ligand, an approach that forms the cornerstone of endocrine therapy in postmenopausal patients (5).

Despite the initial efficacy of these agents, the acquisition of therapeutic resistance remains a pervasive clinical challenge. While *in-vitro* reprogramming of cell lines serves as the predominant model for elucidating resistance (6,7), there remains a paucity of robust data comprehensively comparing the mechanistic divergence between acute therapeutic response and established, long-term resistance. The eventual evolution of resistance is rooted in both non-genetic reprogramming and loss of estrogen receptor expression (8) as well as via acquisition of genetic mutations in the Estrogen Receptor itself (9–11). We posit that the biological manifestation of resistance is intrinsically defined by the specific modality of receptor or biosynthetic pathway inhibition as SERMs like tamoxifen achieve partial loss of ERα compared to SERDs like fulvestrant which degrade the receptor (12). The evolution of therapeutic resistance is thus likely to involve a compendium of molecular adaptations contingent on therapy modality. In its earliest phases, acute drug exposure often triggers transient compensatory mechanisms, allowing a subpopulation of “persister” cells to survive the initial pharmacological onslaught across various types of cancer, including ER+ breast cancer (13,14). However, under the sustained pressure of chronic exposure, these temporary adaptations can transition into a more permanent, “locked-in” resistant state accompanied by profound alterations in the epigenetic landscape (7,15). Parallel to these non-genetic shifts, the acquisition of somatic mutations, most notably the Y537S and D538G variants within the ERα protein, grants cancer cells constitutive, ligand-independent signaling capabilities especially under the influence of Aromatase Inhibitor treatments (9–11).

The cumulative effect of these genetic and epigenetic disruptions can result in a massive transcriptomics reprogramming that extends beyond simple oncogenic receptor signaling (16–18). This remodeling can be conceptualized as a dual assault on cellular identity; it not only recalibrates the oncogenic signaling pathways upon which the untreated cells once depended, but also fundamentally shifts the cellular state to explore a phenotypic “possibility space” of favorable survival mechanisms. This plasticity can include the loss of tissue- and lineage-specific markers, as well as the gain of undifferentiated and epithelial-mesenchymal transition (EMT) gene programs that directly enhance metastatic potential (18–24). Furthermore, such large-scale rewiring of the transcriptome inevitably alters other associated hallmarks of cancer such as the cancer cells’ immunogenicity, modulating the expression of immune-related genes that dictate visibility to the host immune system (25–28). Despite the clinical gravity of these adaptations, a systematic understanding of how specific endocrine therapeutic modalities differentially sculpt these transcriptomic landscapes remains a critical void in the current literature.

Given emerging links between lineage plasticity, partial epithelial–mesenchymal transition (pEMT/ hybrid E/M), and therapy adaptation, it remains unclear whether distinct endocrine therapies channel tumor evolution along shared or divergent reprogramming trajectories. In particular, the extent to which the specific mode of ER targeting shapes lineage identity and epithelial–mesenchymal state has not been systematically examined. Here, we integrate multi-omic profiling across SERM, SERD, and ESR1-mutant contexts to define how drug-specific selective pressures sculpt the adaptive landscape of ER+ breast cancer by facilitating both lineage and epithelial-mesenchymal reprogramming with immunological and microenvironmental altering consequences.

## Results

### *In-vitro* Trajectories of Endocrine Resistance Defined by Treatment Duration and Drug Modality

*In-vitro* model systems have been instrumental in elucidating the molecular underpinnings of ET response, providing a controlled environment to dissect the complex ERα signaling (6). However, there is a notable scarcity of studies that perform a direct comparison across independent datasets to evaluate how the transcriptional states induced by SERMs, SERDs, and ESR1 mutations converge or diverge. A primary obstacle to achieving this objective has been the pervasive batch effects, in which technical variation, arising from differences in cell line passage, media composition, and sequencing platforms, often masks the underlying biological signal. Without such cross-study comparisons, it remains difficult to determine which transcriptomic shifts represent robust, recurring transcriptional programs.

To address this gap, we compiled 27 publicly available transcriptomic datasets focused on the MCF7 ER+ breast cancer cell line (**Supplementary Table 1**), focusing on both control and treated MCF7 populations capturing the acute and chronic transcriptional responses to short-term (minutes to days) and long-term exposure (months) to tamoxifen (SERM) and fulvestrant (SERD) (**Supplementary Table 1**). We specifically included estradiol (E2)-stimulated controls alongside ESR1 (Y537S) mutant lines to contrast ligand-dependent versus constitutive, ligand-independent receptor signaling, a frequent resistance mechanism after prolonged treatment with Aromatase Inhibitors (9). Furthermore, we incorporated 10 basal-like breast cancer cell lines to serve as a biological outgroup, allowing us to clearly distinguish therapy-induced transcriptional changes from fundamental differences in disease subtype. By aggregating these disparate sources, we aimed to create a robust baseline for comparing primary therapeutic responses against the divergent pathways that emerge during long-term resistance (***Figure 1A***).

**Figure 1:**
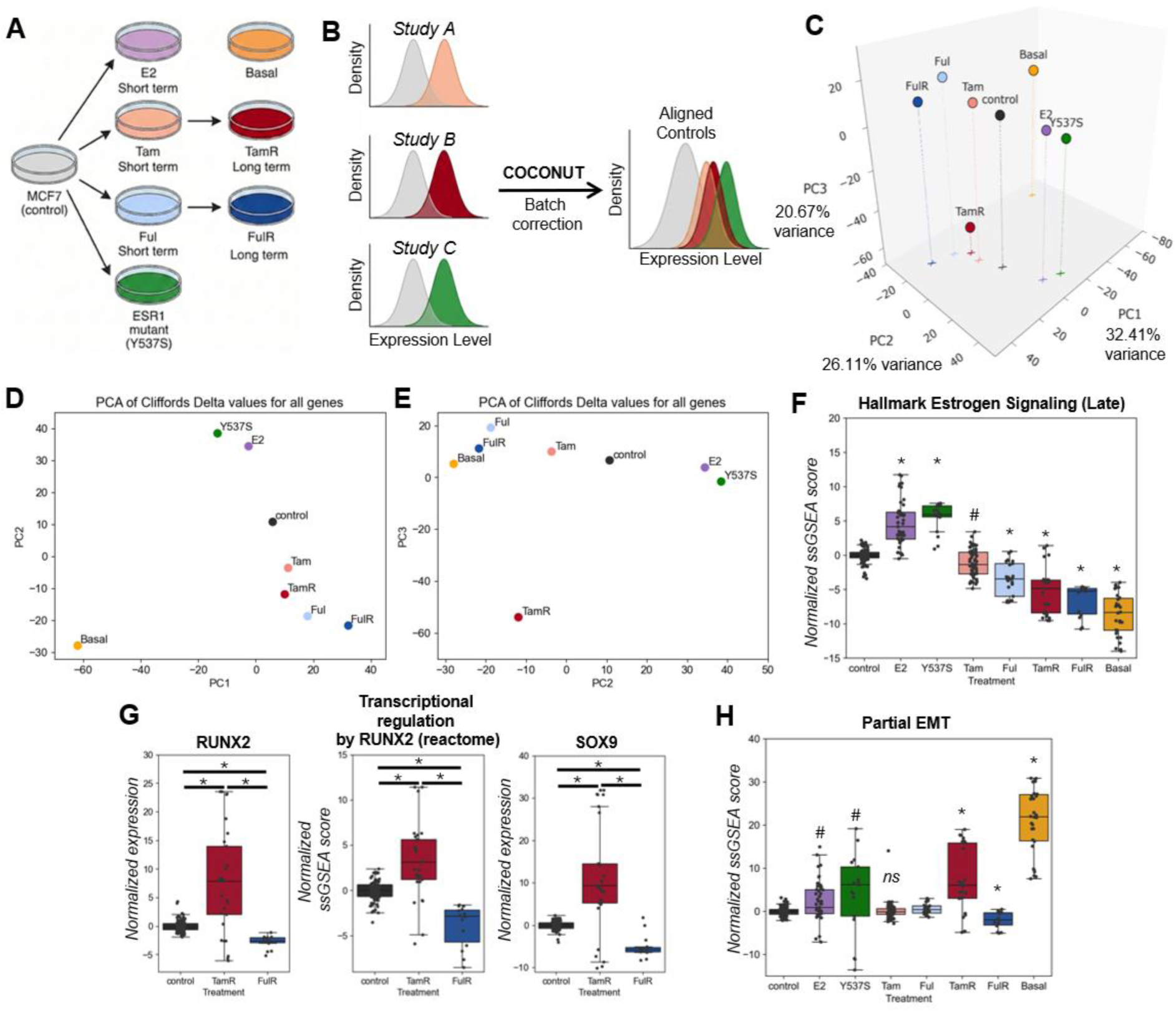
Distinct transcriptional trajectories across modes of endocrine perturbation. (A) Schematic overview of the in vitro experimental models used for comparative analysis, including control MCF7, estradiol treated (E2), tamoxifen-treated (Tam), tamoxifen-resistant (TamR), fulvestrant-treated (Ful), fulvestrant-resistant (FulR), ESR1-mutant (Y537S) conditions, and a panel of Basal breast cancer cell lines serving as a biological outgroup. (B) Diagram illustrating cross-study batch correction performed using COCONUT to harmonize gene expression distributions across datasets prior to comparative analysis. (C) Three-dimensional principal component analysis (PCA) plot generated from Cliff’s delta effect sizes for each gene across conditions, demonstrating the relative positioning of experimental groups within transcriptional space. (D) Two-dimensional PCA projection of PC1 versus PC2 derived from the same Cliff’s delta matrix. (E) Two-dimensional PCA projection of PC2 versus PC3. (F) Boxplots depicting normalized ssGSEA scores for the Hallmark Estrogen Signaling (Late) gene set across conditions (Gene signature source: MSigDB). (G) Boxplots showing (i) normalized RUNX2 expression, (ii) ssGSEA scores for the Reactome RUNX2 transcriptional regulation pathway (Gene signature source: MSigDB), and (iii) normalized SOX9 expression across control MCF7, TamR, and FulR conditions. (H) Boxplots displaying normalized ssGSEA scores for the partial epithelial–mesenchymal transition (pEMT) gene set across conditions (Gene signature source: Puram et al. (2017)). For all boxplots, * denotes a large effect size with a significant p-value (<0.01, Mann–Whitney U test) compared to control MCF7 cells; # indicates a medium effect size with p < 0.01; ns denotes either a small effect size or a non-significant difference relative to control.

To correct for the technical variability between different studies, we applied the COmbat CO-Normalization Using conTrols (COCONUT) algorithm (29) to batch-correct the data. COCONUT aligns the control samples from each study to a common baseline and effectively removes technical noise while preserving the biological effects on the treatment groups (***Figure 1B***). To quantify transcriptional changes across therapeutic conditions, we calculated effect sizes for each gene with respect to control MCF7 cells using Cliff’s delta, a non-parametric measure robust to non-normal distributions and unequal sample sizes while statistical significance was assessed using the Mann–Whitney U test (**See Methods**). Principal component analysis (PCA) of the Cliff’s delta values for all genes across different conditions compared to their respective control (**Supplementary Data 1**) revealed that ∼80% of cross-sample variation was explained by the first three principal components (**Figure 1C**). The primary axis of variation (PC1; 32.41% variance explained) majorly reflected intrinsic subtype differences, separating basal-like cell lines from the luminal MCF7 model (**Figure 1D**) assessed using breast cancer specific luminal and basal signatures (30) (**Supplementary Figure 1A**). The second principal component (PC2; 26.11% variance explained) segregated short-term endocrine treatments (tamoxifen [Tam] and fulvestrant [Ful]), long-term resistance models (TamR and FulR), and basal cell lines from estradiol (E2)-treated and ESR1-mutant conditions (**Figure 1E**). To further establish the links between PC2 and estrogen receptor (ERα) signaling activity, we computed single-sample gene set enrichment analysis (ssGSEA) scores for Hallmark estrogen response pathways (early and late). As expected, E2-treated and ESR1-mutant models exhibited elevated estrogen signaling activity compared to the basal cell lines (**Figure 1F, Supplementary Figure 1B**). Notably, long-term endocrine-resistant models (TamR and FulR) showed globally reduced ERα signaling. In short-term treatments, fulvestrant induced a more pronounced suppression of ERα signaling than tamoxifen (**Figure 1F, Supplementary Figure 1B**), consistent with its mechanism of action as a selective estrogen receptor degrader (SERD) that promotes rapid ERα degradation and attenuates downstream transcriptional responses (12).

The third principal component (PC3) further distinguished TamR from FulR states (**Figure 1E**), indicating substantial transcriptional divergence between these two endocrine resistance phenotypes. While FulR was primarily characterized by near-complete loss of estrogen signaling, TamR exhibited only partial attenuation of ERα activity and a distinct PC3-associated transcriptional program. We additionally performed Reactome pathway enrichment analysis of genes with high positive or negative PC3 loadings using (**Supplementary Data 2**). IFN-α and IFN-γ signaling pathways were significantly associated with positive PC3 loadings (**Supplementary Data 2**). In contrast, RUNX-mediated transcriptional programs were associated with negative PC3 loadings, highlighting RUNX2 pathway activation specifically in TamR (**Figure 1G, i–ii, Supplementary Data 2**). These findings are consistent with prior reports demonstrating that the ERα–RUNX2 axis contributes to transcriptional reprogramming and adaptive resistance in tamoxifen-resistant breast cancer models (31). Furthermore, SOX9, a downstream target of the ERα–RUNX2 complex (31), was selectively upregulated in TamR samples (**Figure 1G, iii**), providing additional support for activation of this regulatory axis.

Given the established association between epithelial–mesenchymal transition (EMT) and therapeutic resistance, we evaluated partial EMT (pEMT) (32) and Hallmark EMT signatures (33) across conditions. Basal cell lines exhibited the highest enrichment for both pEMT and Hallmark EMT signatures (**Figure 1H, Supplementary Figure 1C**), consistent with previous reports describing basal-like breast cancer models as exhibiting partial EMT features. Notably, ESR1-Y537S mutant models, TamR, and E2-treated conditions also demonstrated significant modulation of pEMT signatures (**Figure 1H**), suggesting that diverse endocrine perturbations, including ligand stimulation, receptor mutation, and acquired resistance, can reshape epithelial–mesenchymal cell states consistent with prior studies (21,34,35).

Collectively, these findings indicate that non-genetic endocrine resistance is broadly characterized by attenuation of canonical estrogen receptor (ERα) signaling. However, resistance arising under selective pressure from SERMs versus SERDs do not converge toward a singular resistant phenotype. Instead, these treatment modalities drive bifurcation into distinct transcriptional states, reflecting drug-specific mechanisms of action and adaptive rewiring of downstream signaling networks.

**Supplementary Figure S1:**
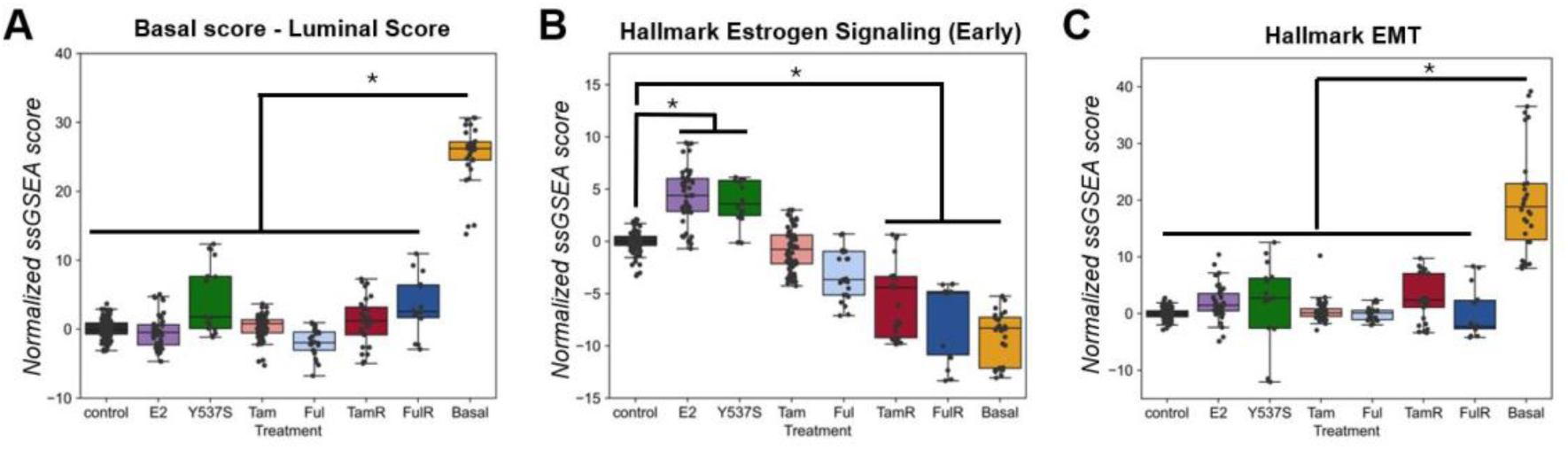
(A) Boxplots displaying normalized ssGSEA scores for an overall basal reprogramming score (basal - luminal) gene set across conditions (Gene signature source: Nair et al. (2024)). (B) Boxplots depicting normalized ssGSEA scores for the Hallmark Estrogen Signaling (Early) gene set across conditions (Gene signature source: MSigDB). (C) Boxplots depicting normalized ssGSEA scores for the Hallmark Epithelial Mesenchymal Transition gene set across conditions (Gene signature source: MSigDB).

### Long-term SERM resistance is associated with partial luminal-to-basal reprogramming and multi-lineage transcriptional mimicry

Lineage plasticity has increasingly been implicated in endocrine resistance in ER+ breast cancer. Notably, most well-characterized examples of lineage reprogramming have been described in the context of ESR1-mutant tumors, intrinsically ER-low or luminal-low subtypes, or prolonged estrogen deprivation models (LTED models) (19,20,25). In contrast, the extent to which prolonged treatment with selective estrogen receptor modulators (SERMs) versus selective estrogen receptor degraders (SERDs) induces comparable lineage remodeling in non-genetic resistance settings remains incompletely defined. To address this question, we quantified pathway activity scores for 6 curated luminal and basal gene signatures across control MCF7 cells, tamoxifen-resistant (TamR), fulvestrant-resistant (FulR), and established basal-like breast cancer cell lines.

As expected, basal cell lines exhibited significantly higher basal signature scores and lower luminal signature scores relative to control MCF7 cells across two independent lineage gene sets (36,37) (**Figure 2A-B**), validating the discriminatory capacity of the signatures used. TamR cells demonstrated a concordant shift, characterized by increased basal and decreased luminal pathway activity, although the magnitude of this shift was substantially smaller than that observed in bona fide basal cell lines (**Figure 2A-B**). In contrast, FulR cells displayed minimal or no significant alterations in lineage-associated gene expression programs relative to control luminal cells (**Figure 2A-B**). These findings indicate that non-genetic endocrine resistance is accompanied by lineage reprogramming to varying degrees, with SERM resistance inducing a more pronounced luminal-to-basal drift than SERD resistance. To further contextualize this shift along mammary developmental trajectories, we quantified enrichment for gene signatures derived from single-cell transcriptomic profiles of mammary stem-like and mature myoepithelial (basal) cell populations (38). TamR cells are preferentially enriched for mature basal programs rather than a full mammary stem-like transcriptional state, whereas basal cancer cell lines were enriched for both developmental programs (**Figure 2C**).

**Figure 2:**
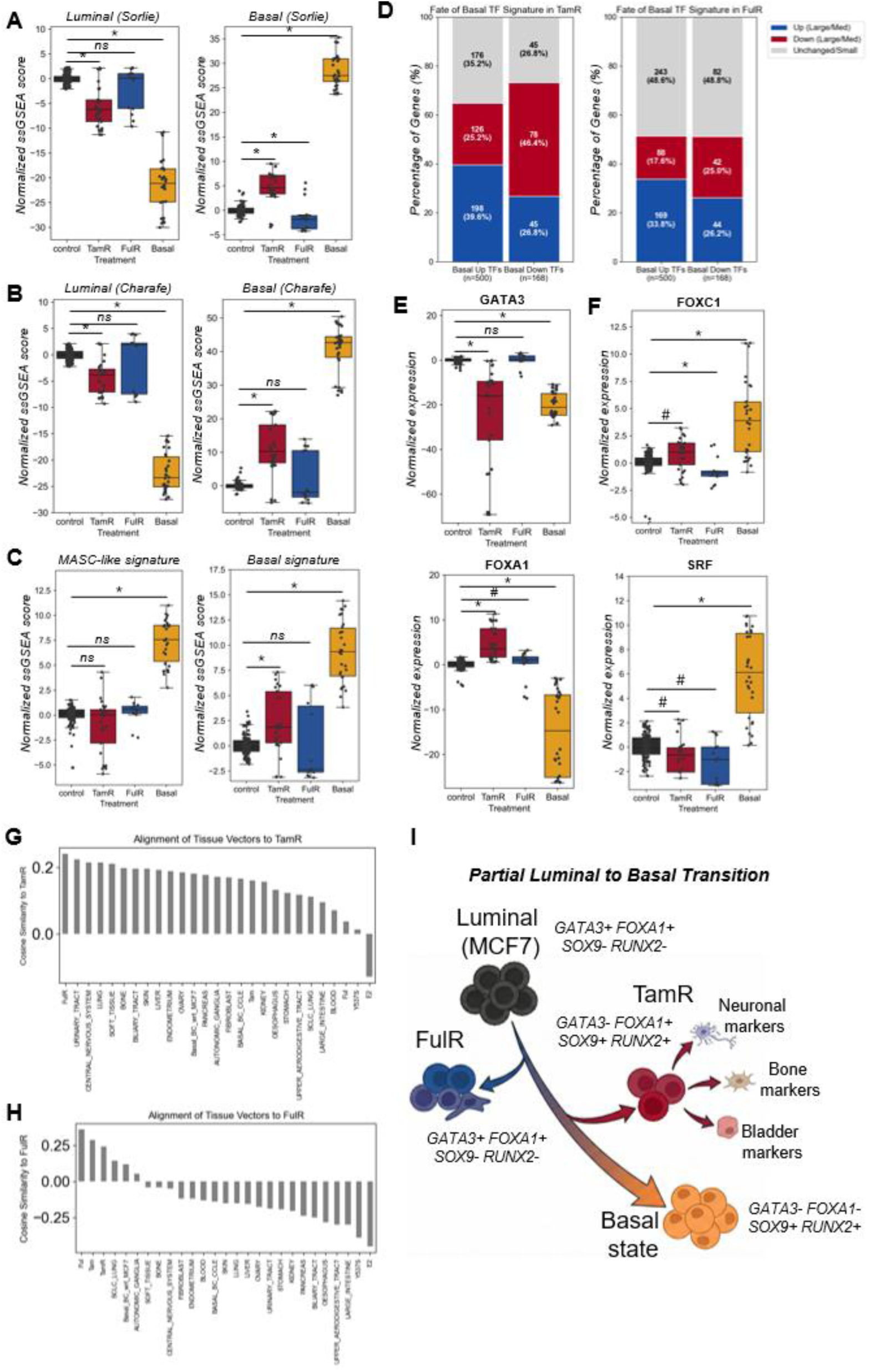
Differential lineage reprogramming and transcription factor remodeling across endocrine resistance states. (A) Boxplots showing normalized ssGSEA scores for the (i) Sorlie Luminal and (ii) Sorlie Basal gene signatures (Gene signature source: Sorlie et al (2003)) across experimental conditions. (B) Boxplots showing normalized ssGSEA scores for the (i) Charafe Luminal and (ii) Charafe Basal gene signatures (Gene signature source: MSigDB). (C) Boxplots displaying normalized ssGSEA scores for the (i) mammary stem cell–like (MASC-like) signature and (ii) single-cell–derived Basal signature (Gene signature source: Saeki et al. (2021)). (D) Stacked bar plots depicting the fraction of transcription factors (TFs) significantly dysregulated in (i) TamR and (ii) FulR relative to the total set of TFs significantly dysregulated in Basal cell lines compared to control MCF7 cells. Red indicates large or medium effect size with significant downregulation; blue indicates large or medium effect size with significant upregulation; grey denotes small effect size or non-significant dysregulation. (E) Boxplots showing normalized expression of (i) GATA3 and (ii) FOXA1 across conditions. (F) Boxplots showing normalized expression of (i) FOXC1 and (ii) SRF across conditions. (G) Cosine similarity analysis comparing treatment conditions and CCLE cell lines (stratified by tissue type) to TamR, using control MCF7 (for treatment comparisons) or luminal cell lines (for CCLE comparisons) as reference baselines. (H) Cosine similarity analysis comparing treatment conditions and CCLE cell lines (stratified by tissue type) to FulR, using control MCF7 or luminal cell lines as reference baselines. (I) Schematic summarizing the extent of partial luminal loss and basal gain in TamR and FulR relative to control MCF7 cells. TamR additionally exhibits activation of multi-lineage programs, including neuronal, bone, and bladder-associated markers. * denotes a large effect size with a significant p-value (<0.01, Mann–Whitney U test) compared to control MCF7 cells; # indicates a medium effect size with p < 0.01; ns denotes either a small effect size or a non-significant difference relative to control.

To interrogate the regulatory basis of this partial reprogramming, we examined differential expression of transcription factors (TFs) across conditions. We first identified TFs significantly altered in basal cell lines relative to MCF7 cells. In total, 500 TFs were significantly upregulated and 168 were significantly downregulated in basal compared to luminal cells. In TamR samples, only 198 of the 500 basal-upregulated TFs (39.6%) were upregulated in the same direction, and 78 of the 168 basal-downregulated TFs (46.4%) were concordantly downregulated (**Figure 2D, i**). Notably, 126 of 500 (25%) TFs expected to increase in a basal transition were instead significantly downregulated in TamR, and 45 of 168 (27%) TFs expected to decrease were paradoxically upregulated (**Figure 2D, i**). These discordant shifts provide strong evidence that TamR cells do not undergo a complete lineage conversion to a basal state but instead adopt a hybrid or intermediate transcriptional identity. FulR samples exhibited a similar but attenuated pattern of remodeling; however, nearly 49% of lineage-associated TFs were not significantly altered in the direction of a basal state (**Figure 2D, ii**), consistent with a more limited degree of lineage plasticity under SERD resistance (**Figure 2A-B**). GATA3 and FOXA1, central regulators of luminal breast epithelial identity (39,40), display divergent behavior across conditions. GATA3 was markedly downregulated in TamR and in basal cell lines but remained largely unchanged in FulR (**Figure 2E, i**). FOXA1 exhibited significant upregulation in TamR, whereas it was downregulated in basal cell lines as expected (**Figure 2E, ii**), suggesting a partial rather than a complete luminal loss. Conversely, basal-associated TFs such as FOXC1 (41) and SRF (42) were robustly upregulated in basal cell lines but showed only modest or negligible changes in TamR and FulR (**Figure 2F, i–ii**), reinforcing the concept of incomplete basal specification. TP63, encoding the ΔNp63 isoform, a key regulator of basal partial epithelial-mesenchymal (EM) identity (43), did not demonstrate large differential expression at the bulk RNA level in either endocrine-resistant or basal cell lines (**Supplementary Figure 2A, i**). However, ΔNp63 pathway activity was significantly enriched in basal cell lines, suggesting isoform-specific regulation or post-transcriptional control not fully captured at total TP63 transcript levels. Importantly, no significant ΔNp63 pathway activation was observed in TamR, and FulR demonstrated pathway downregulation (**Supplementary Figure 2A, ii**), further supporting the conclusion that endocrine resistance does not culminate in a bona fide basal lineage state.

Functional enrichment analysis among genes associated with the TamR-specific PC3 program revealed enrichment of developmental processes including osteoblast development and differentiation, chondrocyte differentiation, forebrain development, and embryonic organ morphogenesis (**Supplementary Data 2**). These findings raised the possibility that TamR cells adopt transcriptional features reminiscent of non-mammary tissues. To formally evaluate cross-lineage similarity, we computed cosine similarity between Cliff’s delta profiles of TamR and FulR samples compared to control MCF7 cells and effect size profiles from diverse cancer cell lines compared to corresponding luminal breast cancer cell lines in the Cancer Cell Line Encyclopedia (CCLE), using luminal breast cancer lines as the reference baseline (**see Methods**). Strikingly, TamR transcriptional profiles demonstrated highest similarity to cancer cell lines derived from bladder, central nervous system (CNS), lung, and bone tissues (**Figure 2G**). In contrast, FulR profiles did not exhibit strong or consistent similarity to specific non-breast tissue lineages (**Figure 2H**). Consistent with these observations, lineage-restricted TFs characteristic of these tissues was selectively upregulated in TamR. The luminal bladder-associated TF PPARG was significantly elevated in TamR (**Supplementary Figure 2B**) (44,45). Similarly, bone and cartilage master regulators RUNX2 and SOX9 (46,47), were upregulated in TamR but not in FulR (**Figure 1G**). Additionally, CNS-associated TFs, including SOX2, NEUROG2, POU2F3, MEIS1, and ISL2 (48–51), were selectively enriched in TamR with minimal effect size changes in FulR or basal cell lines (**Supplementary Figure 2C**). Collectively, these data indicate that TamR cells acquire a composite transcriptional state characterized by partial basal reprogramming coupled with activation of multi-tissue developmental programs.

**Supplementary Figure S2:**
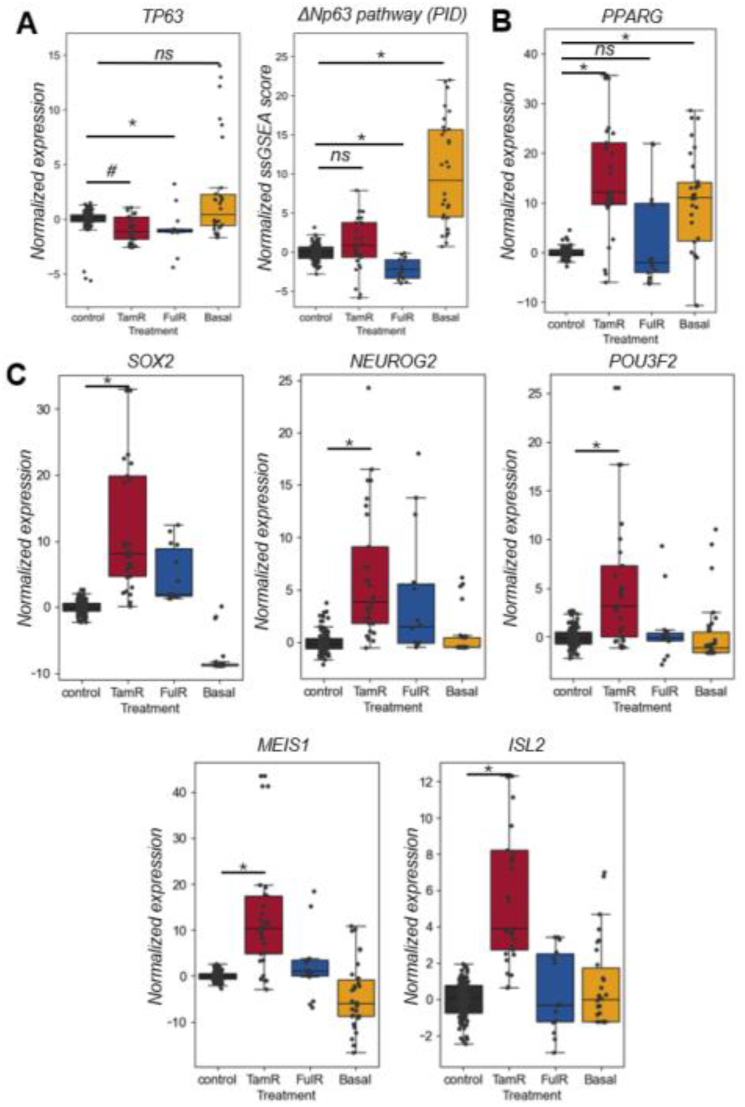
**(A)** Boxplots showing (i) normalized TP63 expression, (ii) ssGSEA scores for the PID ΔNp63 pathway (Gene signature source: MSigDB) **(B)** Boxplots showing normalized expression of PPARG across control, TamR, FulR and Basal conditions. **(C)** Boxplots showing normalized expression of brain specific transcription factors SOX2, NEUROG2, POU2F3, MEIS1, and ISL2 across conditions. * denotes a large effect size with a significant p-value (<0.01, Mann–Whitney U test) compared to control MCF7 cells; # indicates a medium effect size with p < 0.01; ns denotes either a small effect size or a non-significant difference relative to control.

Taken together, these results demonstrate that long-term tamoxifen resistance induces a discernible, yet incomplete, luminal-to-basal lineage drift accompanied by transcriptional mimicry of multiple non-mammary tissue programs. In contrast, fulvestrant resistance is associated with substantially weaker lineage remodeling and limited cross-lineage activation. These findings highlight that distinct endocrine therapies impose unique selective pressures, resulting in divergent adaptive trajectories rather than a unified resistant endpoint.

### Lineage reprogramming in SERM resistance is characterized by stabilization of a partial epithelial–mesenchymal state linked to acquired basal identity

Lineage plasticity in endocrine resistance has traditionally been conceptualized as a binary transdifferentiation process in which luminal identity is relinquished in favor of a basal-like state (19,20,25). In parallel, numerous studies have independently associated epithelial–mesenchymal transition (EMT) features with endocrine resistance (21,52). However, phenotypic heterogeneity in breast cancer arises from the intersection of two partially coupled biological axes: (i) a tissue-lineage axis (luminal versus basal) and (ii) a morphological axis (epithelial versus mesenchymal) (53). Although luminal breast cancers typically maintain a relatively stable epithelial phenotype, multi-omic profiling studies have demonstrated that basal-like breast cancer is intrinsically associated with a hybrid epithelial/mesenchymal (E/M) state rather than a fully mesenchymal identity (53). This hybrid state is often described as partial EMT (pEMT), reflecting coordinated but incomplete activation of mesenchymal programs alongside retention of epithelial features. Building on our observation that TamR transcriptomes exhibit only partial concordance with canonical basal lineage programs (**Figure 2A-C**), we hypothesized that SERM-induced resistance does not drive a complete EMT inversion but instead stabilizes a distinct partial E/M phenotype that is mechanistically linked to basal-like transcriptional regulation.

To formally characterize this “partial” reprogramming, we deconvolved transcriptional dynamics across three regulatory layers: pan-cancer epithelial transcription factors (TFs), pan-cancer mesenchymal TFs, and breast-specific ERα-associated TFs (**see Methods**). Using the Cancer Cell Line Encyclopedia (CCLE) compendium, we identified 73 pan-cancer epithelial TFs defined as TFs positively correlated (Spearman ρ > 0.4) with a cancer agonistic epithelial signature (54) in at least 6 of 23 solid tumor tissue types, thereby minimizing tissue-specific outliers. Similarly, 87 mesenchymal TFs were defined using the same cross-tissue criteria against a mesenchymal gene signature (54). To isolate lineage-restricted oncogenic signaling specific to breast cancer, we further defined a set of 47 breast-specific ERα signaling-associated TFs. These TFs showed strong correlation (ρ > 0.3) with the Hallmark Estrogen Response pathway specifically within breast cancer cell lines and exhibited a Z-score > 1.5 relative to their correlation distribution across all tissue types (**Figure 3A**). This filtering ensured enrichment for TFs governing ERα-driven transcriptional programs rather than generic epithelial identity. Importantly, overlap between the pan-cancer epithelial TFs and breast-specific ERα-associated TFs was minimal (3 genes; Jaccard index = 0.025), demonstrating that they represent largely independent regulatory modules. Across all breast cancer lines in CCLE, epithelial and mesenchymal TF scores were strongly negatively correlated (r = −0.78, p < 0.01), while epithelial and ERα-associated TFs were positively correlated (r = 0.58, p < 0.01) (**Supplementary Figure S3A, i-ii**). Mesenchymal TF scores were negatively correlated with ERα-associated TFs (r = −0.78, p < 0.01) (**Supplementary Figure S3A, iii**). Additionally, within each TF category, members exhibited strong positive intra-group correlations and reciprocal negative correlations across groups (**Supplementary Figure S3B**), consistent with coordinated regulatory “teams”.

**Figure 3:**
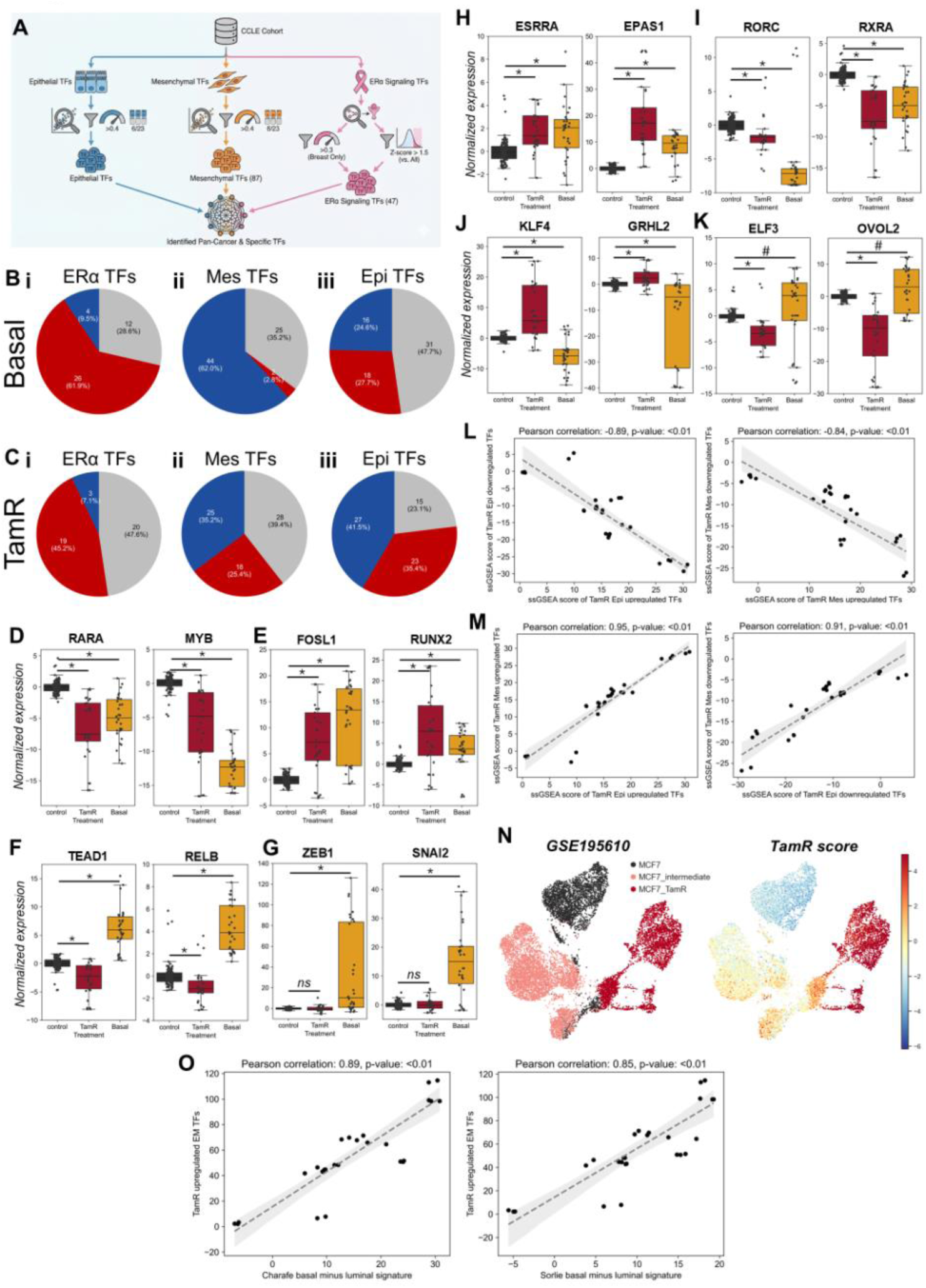
Coordinated epithelial–mesenchymal transcription factor reprogramming defines the TamR partial EM state. (A) Schematic illustrating the identification and classification of pan-cancer epithelial and mesenchymal transcription factors (TFs), along with breast cancer–specific ERα-associated TFs, derived from CCLE cell line datasets. (B) Pie charts showing the fraction of (i) ERα-associated TFs, (ii) mesenchymal TFs, and (iii) epithelial TFs that are significantly upregulated (blue) or downregulated (red) in Basal cell lines relative to control MCF7 cells. (C) Pie charts showing the fraction of (i) ERα-associated TFs, (ii) mesenchymal TFs, and (iii) epithelial TFs significantly upregulated (blue) or downregulated (red) in TamR relative to control MCF7 cells. Boxplots showing normalized expression across conditions for: (D) ERα-associated TFs RARA and MYB; (E) mesenchymal TFs FOSL1 and RUNX2; (F) mesenchymal TFs TEAD1 and RELB; (G) mesenchymal TFs ZEB1 and SNAI2; (H) epithelial TFs ESRRA and EPAS1; (I) epithelial TFs RORC and RXRA; (J) epithelial TFs KLF4 and GRHL2; (K) epithelial TFs ELF3 and OVOL2. (L) Scatter plots with linear regression lines showing correlations within TamR samples between ssGSEA scores of TamR-upregulated epithelial TFs and TamR-downregulated epithelial TFs, and similarly between upregulated and downregulated mesenchymal TF modules. (M) Scatter plots with regression lines showing correlations in TamR samples between TamR-upregulated epithelial TFs and TamR-upregulated mesenchymal TFs, as well as between TamR-downregulated epithelial and mesenchymal TF modules. (N) UMAP projections of single-cell RNA-seq data depicting parental MCF7 cells, intermediate time-point tamoxifen-treated cells, and fully resistant TamR cells. Cells are additionally colored by the TamR composite score (epithelial upregulated + mesenchymal upregulated – epithelial downregulated – mesenchymal downregulated). (O) Scatter plots showing correlations between TamR-upregulated TF modules (epithelial upregulated + mesenchymal upregulated) and basal–luminal gradients derived from Charafe and Sorlie gene signatures (Basal minus Luminal ssGSEA scores). * denotes a large effect size with a significant p-value (<0.01, Mann–Whitney U test) compared to control MCF7 cells; # indicates a medium effect size with p < 0.01; ns denotes either a small effect size or a non-significant difference relative to control.

As expected, basal cell lines displayed large-scale suppression of ERα-associated TFs, with 61.9% significantly downregulated relative to control MCF7 cells (**Figure 3B, i**). Concomitantly, 62% of pan-cancer mesenchymal TFs were upregulated (**Figure 3B, ii**). Notably, pan-cancer epithelial TFs exhibited a heterogeneous pattern, with 27.7% downregulated and 24.6% upregulated (**Figure 3B, iii**). This mixed epithelial profile, alongside mesenchymal activation and ERα suppression, reinforces the established concept that basal breast cancer represents a partial EM phenotype rather than a complete mesenchymal transition. Thus, basal lineage identity and pEMT are inherently coupled. TamR samples exhibited a more nuanced and fragmented reprogramming landscape. Although, consistent with reduced canonical ERα signaling, 45% of breast-specific ERα TFs were downregulated (**Figure 3C, i**), the modulation of epithelial and mesenchymal TFs was strikingly bifurcated. Within both epithelial and mesenchymal TF lists, nearly equal proportions of genes were upregulated and downregulated (**Figure 3C, ii–iii**), in contrast to only the epithelial program showing a coordinated change in expression in basal cell lines. This regulatory “fractionation” indicates that TamR cells do not undergo a binary epithelial-to-mesenchymal switch. Instead, they stabilize a selective hybrid configuration characterized by simultaneous retention of specific epithelial determinants and acquisition of defined mesenchymal regulators. Importantly, this hybrid state is similar to, but does not fully recapitulate, the pEMT architecture intrinsic to basal-like breast cancer, suggesting partial convergence along the lineage–morphology axis.

Breast cancer–specific ERα target genes such as MYB and RARA were significantly downregulated in both TamR and basal cell lines, confirming attenuation of canonical ERα signaling (**Figure 3D**). Mesenchymal-associated TFs including FOSL1 and RUNX2 were upregulated in both contexts (**Figure 3E**), reinforcing a shared pEMT component. However, divergence emerged within mesenchymal subsets. TFs such as TEAD1 and RELB were upregulated in basal cell lines but significantly downregulated in TamR (**Figure 3F**), while canonical EMT drivers including SNAI2 and ZEB1 were significantly altered in basal lines but remained unchanged in TamR (**Figure 3G**). Thus, TamR activates only a restricted subset of mesenchymal regulators, preventing full mesenchymal conversion and supporting stabilization of a partial EMT state. Epithelial-associated TFs further illustrated the hybrid nature of TamR. TFs such as EPAS1 and ESRRA were upregulated in both TamR and basal lines (**Figure 3H**), consistent with shared pEMT-linked epithelial remodeling. In contrast, TFs including RORC and RXRA were downregulated in basal lines (**Figure 3I**), reinforcing the partial E/M nature of basal identity. Unexpectedly, distinct epithelial TF subsets displayed reciprocal behavior between TamR and basal states. For example, KLF4 and GRHL2 were upregulated in TamR but downregulated in basal cell lines (**Figure 3J**), whereas another subset was downregulated in TamR but upregulated in basal cells (**Figure 3K**). Collectively, we identified four TF submodules, two epithelial and two mesenchymal, that violated classical self-activation and cross-repression patterns. These submodules were reproducibly correlated across the different COCONUT normalized TamR datasets (**Figure 3L–M**), indicating a conserved, therapy-selected regulatory architecture. We consolidated these four TF modules into composite gene signatures representing TamR-specific epithelial and mesenchymal subprograms. Application of these signatures to an independent single-cell dataset (GSE195610) (55) capturing progression from parental MCF7 cells through intermediate persister states to fully resistant TamR cells demonstrated a graded activation trajectory. Specifically, the TamR-associated pEMT program progressively increased as cells transitioned towards resistance (**Figure 3N, Supplementary Figure S3D-E**). Notably, ERα signaling attenuation was most pronounced in fully reprogrammed TamR cells, with intermediate suppression observed in persister states (**Supplementary Figure S3C-E**). Importantly, established basal lineage signatures (e.g., Sorlie or Charafe-derived basal and luminal gene sets) were positively correlated with the TamR-associated pEMT program (**Figure 3O**), further reinforcing the link between partial EMT activation and basal-like transcriptional features in SERM resistance.

**Supplementary Figure S3:**
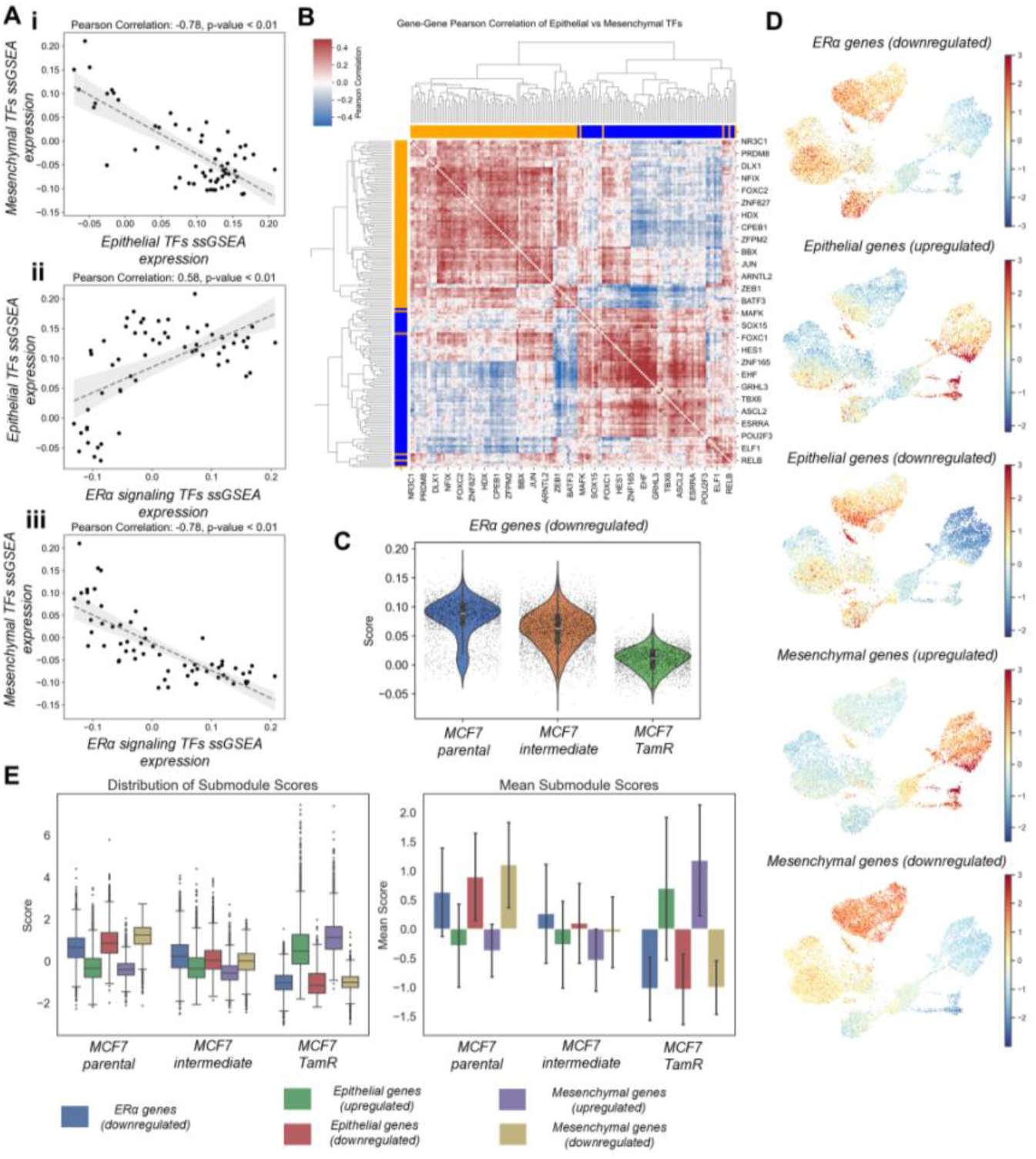
**(A)** Scatterplot showing the correlation between ssGSEA scores of (i) pan-cancer epithelial and mesenchymal TFs, (ii) breast cancer specific ERα TFs and pan-cancer epithelial TFs, and (iii) breast cancer specific ERα TFs and pan-cancer mesenchymal TFs across CCLE breast cancer cell lines. Pearson correlation coefficients and p-values are reported. (B) Hierarchically clustered Gene-gene Pearson correlation matrix between pan-cancer epithelial and mesenchymal TFs across CCLE breast cancer cell lines. Orange and Blue labels indicate Mesenchymal and Epithelial transcription factors respectively. (C) Violin plots showing the expression of breast cancer specific negatively downregulated ERα associated genes in MCF7 parental, MCF7 intermediate and MCF7-TamR single cell RNA seq dataset. (D) Umap plots showing the AUCell assessed pathway activity of submodules (upregulated and downregulated) of ERα associated, Epithelial and Mesenchymal gene sets inferred from bulk COCONUT normalized datasets. (E) Boxplot (left) and bar plot (right) showing the distribution and mean submodule scores for the upregulated and downregulated gene sets belonging to ERα associated, Epithelial and Mesenchymal programs.

Taken together, our analysis reconciles previously disparate observations linking tamoxifen resistance, EMT features, and basal-like transcriptional rewiring. Rather than representing a complete luminal-to-basal transdifferentiation or a full EMT conversion, SERM resistance stabilizes a distinct partial E/M phenotype that is mechanistically and transcriptionally connected to basal identity yet remains molecularly incomplete. This pEMT–basal coupling provides a unifying framework explaining how epithelial-associated regulators (e.g., FOXA1, GRHL2, EPAS1) can coexist with mesenchymal drivers (e.g., SOX9, RUNX2) in the establishment of the TamR state (24,31,56–58).

### Lineage reprogramming in TamR is facilitated by epigenetic reprogramming, increased ERα/FOXA1 binding, and a dysfunctional signaling

Having established that long-term SERM resistance stabilizes a partial epithelial–mesenchymal (pEMT) state that is transcriptionally linked to partial basal reprogramming, we investigated if chromatin organization and epigenetic mechanisms have a role to play in maintaining this intermediate lumino-basal and partial E/M phenotype. We focused on higher-order chromatin architecture using Hi-C dataset generated from parental MCF7, TamR, and FulR cells (GSE118712) (7), with particular attention to key lineage regulators.

Given the marked and specific downregulation of GATA3 in TamR (**Figure 2E**), we first examined chromatin organization at the GATA3 locus. While topologically associating domain (TAD) boundaries (GSE118712) remained largely conserved across MCF7, TamR, and FulR cells, A/B compartment analysis revealed a striking switch at the GATA3 locus in TamR (**Figure 4A**). Specifically, this region transitioned from an active A compartment in MCF7 to an inactive B compartment in TamR, consistent with transcriptional silencing. This compartment switch was accompanied by coordinated loss of ERα, FOXA1, and GATA3 binding across multiple regulatory elements within the affected region (chr10: 77.75–91.25 Mb) (**Figure 4B**). Concomitantly, active enhancer-associated histone marks (H3K27ac) were reduced, indicating functional loss of enhancer activity (**Figure 4B**). Polymer modeling (**see Methods**) of the GATA3 locus (chr10: 3Mb-15Mb) demonstrated enhanced A/B compartment segregation (segregation ratio of 7.54 in MCF7 vs 2.67 in equilibrium control model), highlighting the model’s ability to reproduce biologically relevant chromatin organization (**Supplementary Figure S4A-C**). TamR exhibited a sharp drop in super enhancer contacts, accompanied by increased non-super enhancer interactions compared to both MCF7 and FulR (**Supplementary Figure S4D**). Furthermore, there was a reduction in super enhancer residence time at the promoter of GATA3 (3.07𝜏 in MCF7, 10.37𝜏 in FulR and 2.4𝜏 in TamR) (**Supplementary Figure S4E**), along with loss of key high-frequency enhancer contacts compared to MCF7 scenario (**Supplementary Figure S4F**). Importantly, these architectural alterations were not observed in FulR cells (**Figure 4A; Supplementary Figure S4D-F**), indicating that GATA3 silencing in TamR is mechanistically linked to higher-order chromatin reorganization. Given the central role of GATA3 in maintaining luminal epithelial identity, its epigenetic inactivation provides a structural basis for luminal erosion and partial lineage destabilization in TamR.

**Figure 4:**
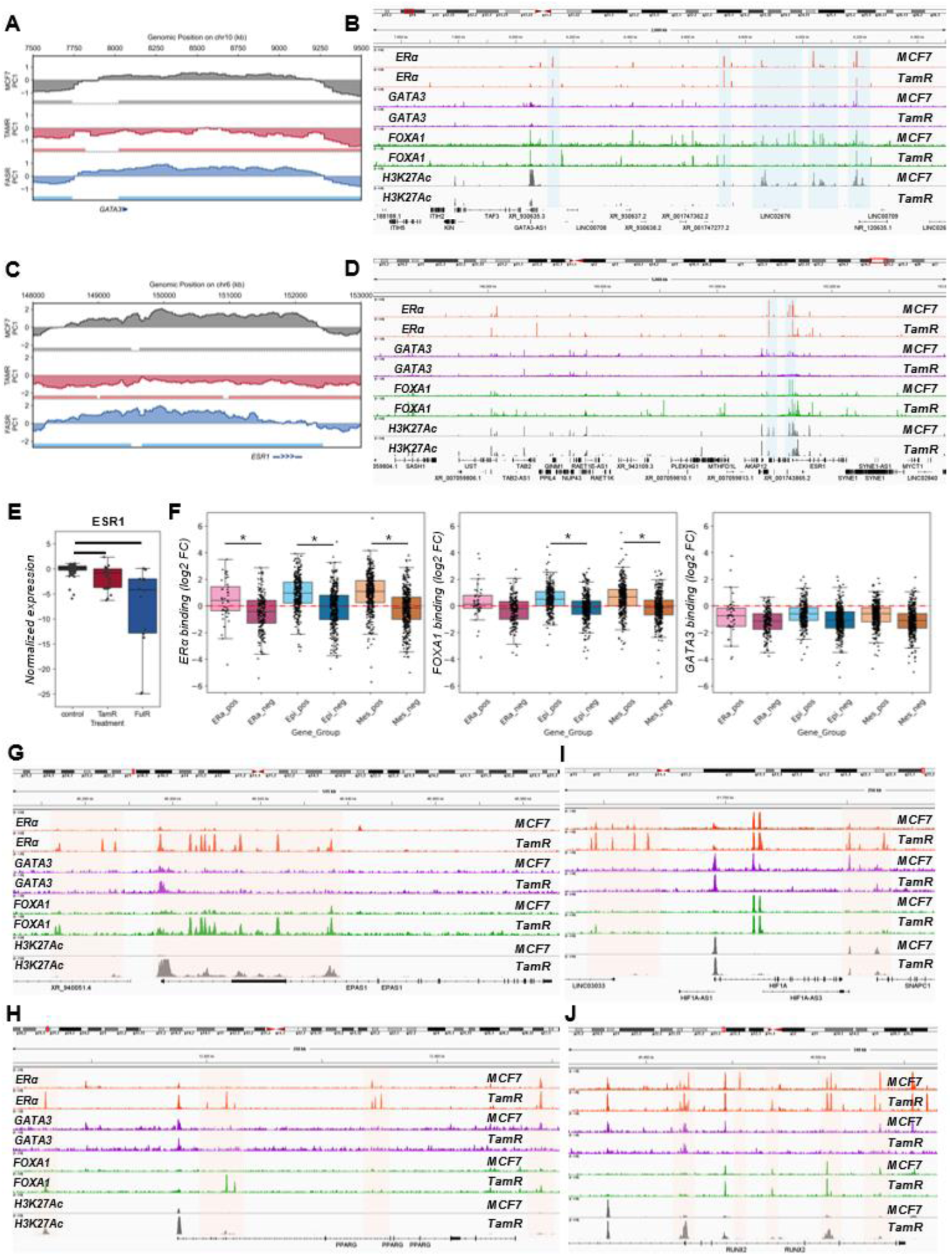
Chromatin compartment switching and ERα cistrome redistribution in TamR cells. (A) A/B compartment eigenvector profiles across the GATA3 locus comparing control MCF7, TamR and FulR cells, demonstrating compartment switching and altered chromatin state shaded regions (horizontal bars) show Topologically associated domains in the different cell lines. (B) Genome browser tracks at the GATA3 locus showing ERα, GATA3, FOXA1, and H3K27ac occupancy in control MCF7 and TamR cells, illustrating reduced luminal enhancer activity and altered transcription factor binding in TamR. Shaded regions showing qualitative changes (C) A/B compartment eigenvector profiles across the ESR1 locus comparing control MCF7, TamR and FulR cells, highlighting A-to-B compartment transition in TamR. (D) Genome browser tracks at the ESR1 locus displaying ERα, GATA3, FOXA1, and H3K27ac occupancy in control MCF7 and TamR cells, demonstrating loss of canonical luminal regulatory architecture in the different cell lines. (E) Boxplot showing normalized ESR1 expression across control MCF7, TamR, and FulR conditions. * denotes significant differences (p < 0.01). (F) Boxplots displaying differential (i) ERα, (ii) FOXA1, and (iii) GATA3 binding in TamR samples (log2 fold change) across defined gene groups with respect to control MCF7 cells, highlighting redistribution of ERα, FOXA1 and GATA3 binding occupancy. * denotes significant differences (p < 0.01). (G) Genome browser tracks at representative TamR-gained loci EPAS1 showing increased ERα binding and H3K27ac signal in TamR relative to control MCF7. (H) Genome browser tracks at a luminal-associated locus PPARG demonstrating reduced enhancer activation and transcription factor occupancy in TamR. (I) Genome browser tracks at the HIF1A locus illustrating TamR-specific enhancer activation and redistribution of ERα, FOXA1, and H3K27ac signals. (J) Genome browser tracks at the RUNX2 locus illustrating TamR-specific enhancer activation and redistribution of ERα, FOXA1, and H3K27ac signals.

We next examined chromatin architecture at the ESR1 locus. In TamR, ESR1 similarly exhibited an A-to-B compartment transition accompanied by alterations in local TAD organization relative to MCF7 (**Figure 4C**). In contrast, neither compartment status nor TAD structure was substantially altered in FulR (**Figure 4C**), despite reduced ERα expression in both resistant states. Although ERα transcript levels were reduced in TamR and FulR (with greater suppression in FulR; **Figure 4E**), the absence of compartment switching in FulR suggests that ERα downregulation in this context is mediated primarily through transcription factor–level or post-transcriptional mechanisms rather than higher-order chromatin restructuring. Collectively, these findings indicate that TamR, but not FulR, undergoes broad epigenetic reconfiguration at core luminal lineage loci, reinforcing that SERM resistance represents a distinct epigenetically reprogrammed state.

We next investigated whether epigenetic remodeling also contributes to stabilization of the TamR-specific pEMT phenotype. We quantified ERα, FOXA1, and GATA3 occupancy at ER-binding sites proximal to genes belonging to (i) the ERα signaling module, (ii) epithelial TF subsets, and (iii) mesenchymal TF subsets that were specifically upregulated or downregulated in TamR (**see Methods**). Strikingly, ERα and FOXA1 binding was significantly enriched at regulatory regions near genes upregulated in TamR, including both epithelial and mesenchymal subsets (**Figure 4F, i–ii**). In contrast, genes downregulated in TamR did not exhibit comparable enrichment. Similarly, ERα signaling–associated genes that were transcriptionally reduced exhibited no enrichment in ERα/FOXA1 occupancy at nearby regulatory loci (**Figure 4F, i–ii**). In contrast to ERα and FOXA1, GATA3 binding was globally diminished across all examined gene sets, consistent with its severe transcriptional downregulation silencing in TamR (**Figure 2E**; **Figure 4F**). These results suggest that while GATA3-mediated luminal reinforcement is epigenetically dismantled, ERα and FOXA1 redistribute to alternative regulatory landscapes.

Notably, despite reduced ESR1 transcript levels in TamR relative to MCF7, ERα occupancy was paradoxically increased at multiple loci associated with the partial epithelial and mesenchymal gene subsets. This observation supports a model in which tamoxifen-bound ERα assumes a non-canonical chromatin-binding profile (15), cooperating with elevated FOXA1 to activate or stabilize a selective pEMT–basal transcriptional program. Examples of such rewiring include newly acquired ERα/FOXA1 binding events near epithelial-associated regulators such as EPAS1 and PPARG, in the absence of GATA3 occupancy (**Figure 4G-H**). Similarly, mesenchymal-associated regulators including HIF1A and RUNX2 displayed emergent enhancer activity marked by H3K27ac and increased ERα/FOXA1 binding (**Figure 4I-J**). These newly established regulatory interactions are consistent with enhancer reprogramming due to ERα binding reported previously (7,15).

Collectively, these findings indicate that TamR resistance is maintained by coordinated epigenetic remodeling that dismantles canonical luminal architecture (via GATA3 compartment silencing), reconfigures ESR1 chromatin topology, and redistributes ERα/FOXA1 binding toward a hybrid transcriptional network. This network integrates selective epithelial determinants with a restricted subset of mesenchymal regulators, reinforcing a stabilized partial EMT state that is intrinsically linked to basal-like lineage features but does not culminate in full basal transdifferentiation. Thus, the partial EM phenotype observed in TamR is not merely transcriptional noise but reflects a structurally encoded chromatin state that mechanistically bridges partial EMT activation with luminal-to-basal lineage plasticity.

**Supplementary Figure S4:**
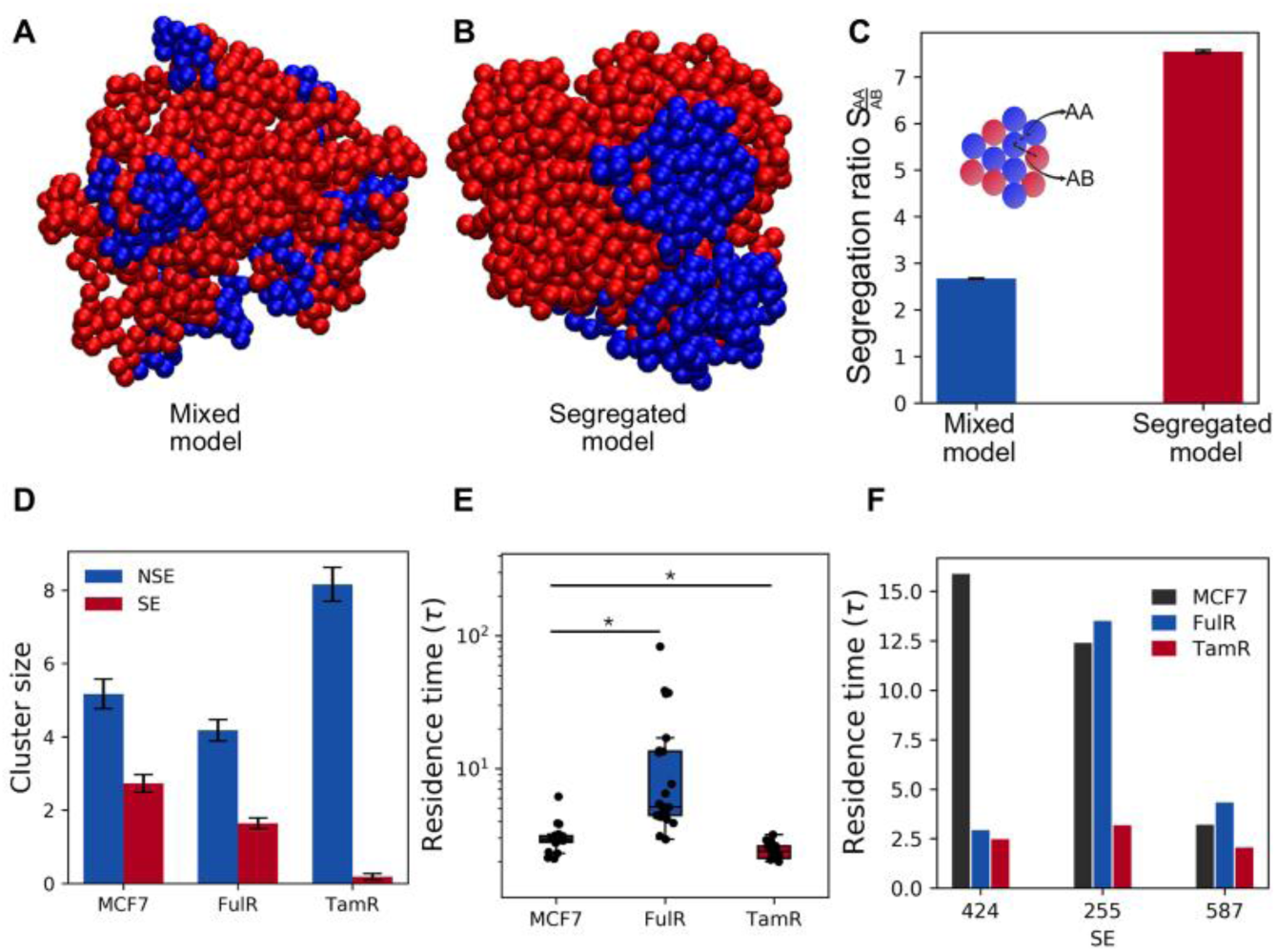
Polymer simulation shows change in super enhancer-promoter interactions due to compartment switching. (A) Polymer snapshot of mixed or equilibrium model without any constraints showing mixing of A compartment (blue beads) and B compartment (red beads), (B) Polymer snapshot of segregated model with constraints showing segregation of A and B compartment of 3 to 15 MB region containing Gata3 gene in MCF7 cell line, (C) Bar plot shows segregation ratio of mixed model and segregated model around Gata3. The error bar shows standard error of mean of segregation ratio. (D) Bar plot shows the cluster size around the Gata3 promoter for non-super enhancers (NSE, blue) and super enhancers (SE, red). Error bars represent the standard deviation of the promoter cluster size across 20 simulations. (E) Box plot shows the distribution of residence time (𝜏) for all super enhancers across the three cell lines. Y-axis is in log scale. Significance was assessed using Mann-Whitney U test. * signifies p-value less than 0.01. p-values of 8.31e-7 (MCF7 vs FulR), 2.54e-3 (MCF7 vs TamR), and 2.99e-7 (FulR vs TamR) (F) Bar plot shows the top three ranked (residence time wise) common super enhancers, with bead indices indicated on the x-axis from left to right.

### Constitutive ESR1 activation does not recapitulate the pEMT–basal reprogramming observed in SERM resistance

To determine whether constitutive estrogen receptor activation phenocopies the transcriptional and epigenetic reprogramming observed in drug-selected resistance, we analyzed ESR1 Y537S mutant models. Both exogenous estrogen (E2) stimulation and activating ESR1 mutations have previously been reported to enhance proliferative signaling and, in some contexts, promote basal cytokeratin expression and partial EMT-associated features (25,34). However, whether this mutational activation converges on the same hybrid pEMT–basal state observed in TamR remains unclear.

Globally, the transcriptional landscape of Y537S mutant cells closely resembled that of acute E2-treated MCF7 samples (**Figure 1C**), with Hallmark Estrogen Response pathway activity reaching levels comparable to E2-stimulated controls (**Figure 1F; Supplementary Figure 1B**). Among 47 transcription factors (TFs) upregulated following E2 treatment, 39 (83%) were also upregulated in the mutant setting. Similarly, of 65 TFs downregulated under E2 stimulation, 34 (52.3%) were concordantly downregulated in Y537S cells, with minimal discordant regulation (**Figure 5A**). These data demonstrate strong concordance between ESR1 mutational activation and ligand-driven ERα signaling, confirming that the mutant state primarily reflects constitutive pathway activation rather than rewiring.

**Figure 5:**
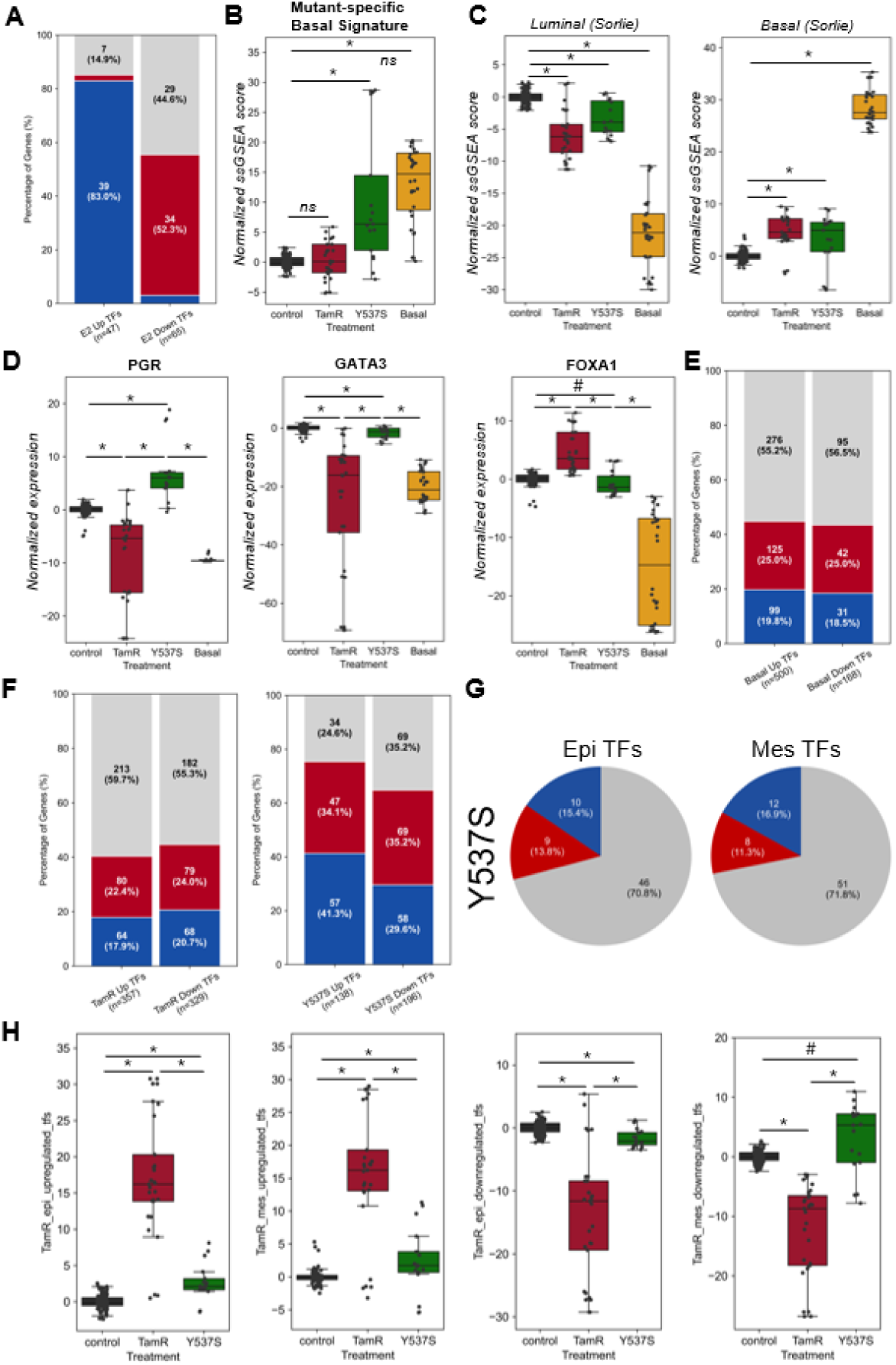
Distinct transcription factor and lineage reprogramming patterns in ESR1 Y537S mutant models. (A) Stacked bar plot showing the fraction of E2-dysregulated transcription factors (TFs) in control MCF7 cells that are significantly dysregulated in MCF7 ERα-Y537S cells. (B) Boxplots showing normalized ssGSEA scores for the mutant-specific basal gene signature. (C) Boxplots displaying normalized ssGSEA scores for the Sørlie Luminal and Sørlie Basal gene signatures across control MCF7, TamR, ERα-Y537S, and Basal cell lines. (D) Boxplots showing normalized expression of PGR, GATA3, and FOXA1 across control MCF7, TamR, ERα-Y537S, and Basal conditions. (E) Stacked bar plot depicting the fraction of mutant-dysregulated TFs that overlap with TFs significantly dysregulated in Basal cell lines relative to control MCF7 cells. (F) Stacked bar plots showing reciprocal overlap between mutant-dysregulated TFs and TamR-dysregulated TFs (i.e., mutant relative to TamR and TamR relative to mutant). (G) Pie charts showing the fraction of pan-epithelial and pan-mesenchymal TFs significantly dysregulated in ERα-Y537S cells, with red indicating downregulation and blue indicating upregulation relative to control MCF7 cells. (H) Boxplots displaying normalized ssGSEA scores for TamR-defined epithelial and mesenchymal subprograms across control MCF7, TamR, and ERα-Y537S conditions. * denotes a large effect size with a significant p-value (<0.01, Mann–Whitney U test) compared to control MCF7 cells; # indicates a medium effect size with p < 0.01; ns denotes either a small effect size or a non-significant difference relative to reference.

Consistent with prior reports linking ESR1 mutations to basal cytokeratin expression and immune pathway activation, we identified upregulation of a mutant-specific basal signature (25) in Y537S cells (**Figure 5B**). This signature was notably absent in TamR samples. Moreover, mutant cells demonstrated significant downregulation of luminal signatures alongside upregulation of basal signatures, with effect sizes comparable to those observed in TamR (**Figure 5C**), indicating a degree of luminal-to-basal drift. However, the regulatory architecture underlying this shift differed substantially from that of TamR. TP63 transcript levels were not significantly altered in mutants, yet ΔNp63 pathway activity was significantly elevated (**Supplementary Figure S5A**), consistent with functional basal pathway engagement. Mechanistically, this divergence appears linked to progesterone receptor (PGR) expression. PGR was significantly upregulated in mutant cells (**Figure 5D, i**), in line with previous studies implicating PGR signaling in mutant-driven reprogramming (25). In contrast, PGR was markedly downregulated in TamR and basal cell lines (**Figure 5D, i**), highlighting a key regulatory distinction between mutational activation and therapy-induced resistance. Although GATA3 was significantly downregulated in mutants (Cliff’s delta: −0.59), the magnitude of repression was considerably less severe than in TamR (Cliff’s delta: −0.93) (**Figure 5D, ii**). A striking dichotomy was observed for the pioneer factor FOXA1: whereas TamR exhibited strong upregulation (Cliff’s delta: 0.92), mutant cells showed significant downregulation (Cliff’s delta: −0.389), with levels comparable to or slightly below parental MCF7 cells (**Figure 5D, iii**). Thus, mutant-driven basal activation occurs in a FOXA1-low, PGR-high context, in contrast to the FOXA1-high/GATA3-silenced architecture that stabilizes the TamR hybrid state.

Analysis of basal lineage TFs further supported the partial and incoherent nature of the mutant transition. Among 500 TFs upregulated in bona fide basal cell lines, 55% were unchanged in mutants; of the remainder, 25% were downregulated and 20% upregulated, without coherent alignment toward a basal state (**Figure 5E**). Similarly, among 168 TFs downregulated in basal lines, 56.5% were unchanged in mutants, with the rest split between up- and downregulation in nearly equal proportions. Concordance between dysregulated TFs in TamR and mutant models was similarly mixed (**Figure 5F**). These findings indicate that although mutant cells activate certain basal-associated features, the transition lacks the coordinated transcriptional restructuring characteristic of basal identity or TamR.

Critically, the epithelial–mesenchymal (EM) axis remained largely stable in mutant cells. Most pan-cancer epithelial and mesenchymal TFs were either unchanged or modestly altered, with substantially less bifurcation than observed in TamR (**Figure 5G**). Quantification of the TamR-specific epithelial and mesenchymal subprograms confirmed that activation of the defined pEMT modules was markedly weaker in mutant cells compared to TamR (**Figure 5H**). Thus, while ESR1 Y537S mutants exhibit partial basal lineage activation, this shift is not tightly coupled to stabilization of a partial E/M state. In contrast to SERM resistance, where pEMT activation and basal drift are mechanistically intertwined, mutational activation primarily reinforces constitutive ERα signaling and PGR-driven programs with only limited remodeling along the EM axis.

Overall, our results indicate that, although both genetic and non-genetic mechanisms can produce elements of luminal erosion and basal activation, only SERM-driven resistance robustly couples basal lineage drift with partial EMT stabilization, underscoring the central role of the pEMT–basal connection in therapy-induced reprogramming.

### The pEMT–basal reprogramming observed in SERM resistance is associated with immune evasion via MHC downregulation and microenvironmental remodeling

Epithelial–mesenchymal plasticity is intricately linked to tumor immunogenicity and immune evasion across diverse solid malignancies (59). While partial EMT (pEMT) states are frequently associated with immunosuppressive phenotypes, often driven by the overexpression of co-inhibitory checkpoints such as CD274 (PD-L1), CD47 and NT5E (CD73) (60–63), the specific immunological consequences of the TamR-associated partial EM state remain undefined. Building upon our observation that interferon (IFN) signaling pathways inversely segregated with the TamR phenotype along principal component 3 (PC3) (**Supplementary Data 2**), we hypothesized that the pEMT-basal transition dictates a specific mode of immune evasion and microenvironmental remodeling. To systematically evaluate this, we first quantified the expression levels of MSigDB Hallmark IFNα and IFNγ pathway genes across our defined samples, TamR and basal cell lines. We observed a significant and coordinated downregulation of both IFNα and IFNγ signaling pathways in TamR samples relative to control MCF7 cells (**Figure 6A**). In stark contrast, the basal cell lines exhibited a robust upregulation of these identical pathways (**Figure 6A**). Furthermore, across both the bulk COCONUT-normalized compendium and our independent single-cell RNA-sequencing trajectories (capturing parental, intermediate, and fully resistant states), the attenuation of IFNα and IFNγ signaling was strongly and negatively correlated with the TamR-specific pEMT composite score (**Figure 6B, Supplementary Figure S5A**). These divergent transcriptomic profiles suggest that while basal-like breast cancers tend to establish an immune-infiltrated or “hot” tumor microenvironment, the partial reprogramming induced by prolonged SERM exposure can drive the tumor toward an immunologically “cold” state probably beyond the treatment naïve immunologically cold luminal tumors (64).

**Figure 6.**
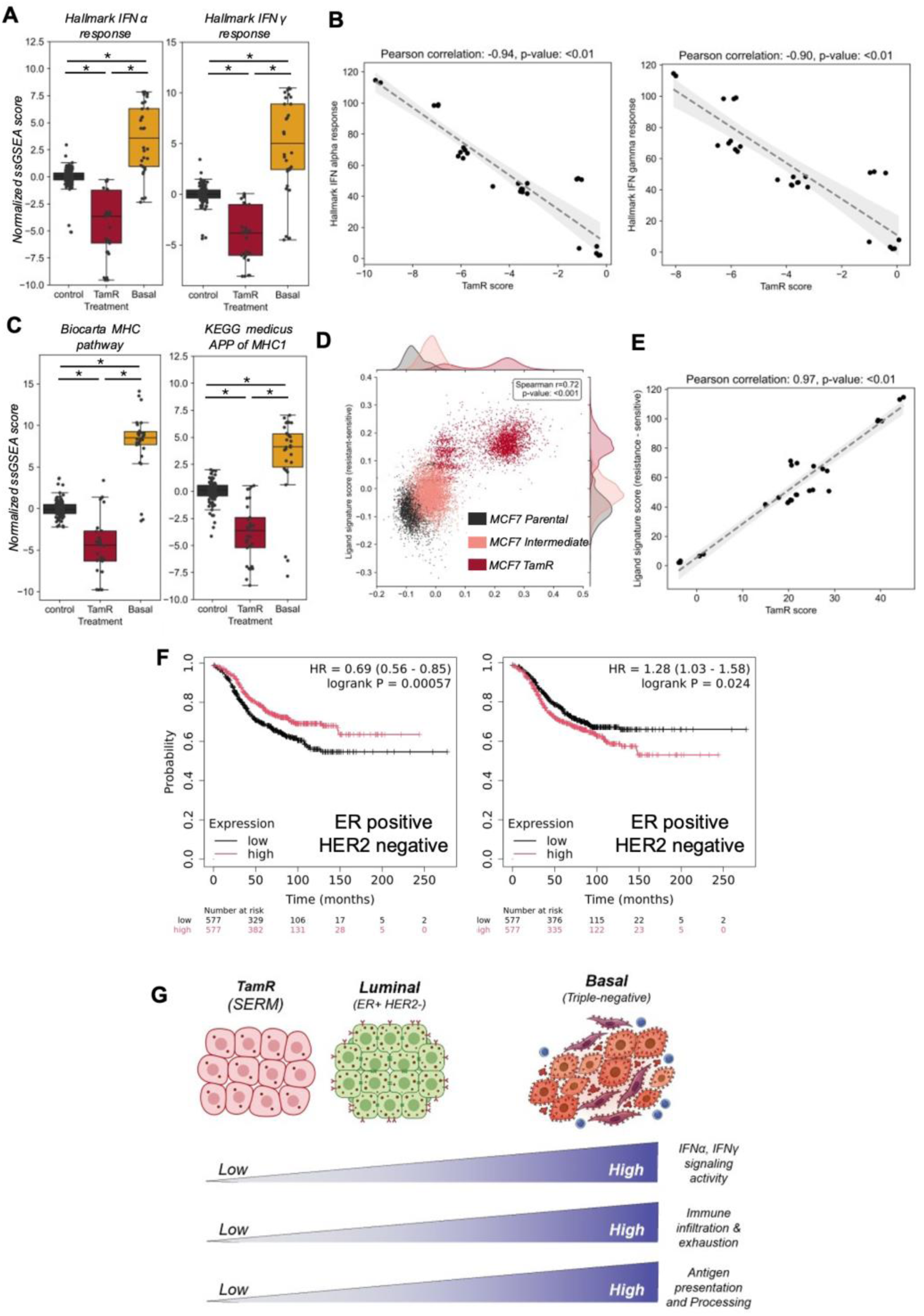
TamR associated pEMT state is associated with suppression of interferon signaling, impaired antigen presentation, and distinct microenvironmental interactions. (A) Normalized ssGSEA scores for Hallmark IFN-α (left) and IFN-γ (right) response pathways across control MCF7, tamoxifen-resistant (TamR), and basal breast cancer cell lines. Statistical significance indicated by * (p < 0.05). (B) Correlation of ssGSEA scores of Hallmark interferon alpha (top) and gamma (bottom) pathway activity with the TamR-pEMT reprogramming score. (C) Boxplots showing ssGSEA scores for antigen presentation pathways, Biocarta MHC pathway (left) and KEGG antigen processing and presentation MHC-1 (right). Statistical significance indicated by * (p < 0.05). (D) Single-cell level correlation between TamR-pEMT score and ligand signature AUCell score (resistant - sensitive). Cells with higher TamR scores show enrichment of the resistant ligand program, with a strong positive Spearman correlation. (E) Bulk-level correlation between TamR-pEMT score and ligand signature score (resistance minus sensitive), demonstrating robust positive association across datasets. (F) Kaplan–Meier survival analysis in luminal breast cancer patients stratified by ligand signatures. Left: high expression of the “sensitive” ligand signature is associated with improved survival (HR = 0.69). Right: high expression of the “resistant” ligand signature correlates with poorer survival (HR = 1.28). (G) Schematic showing downregulation interferon pathways, antigen presentation/processing, suppression of immune cell infiltration and exhaustion in TamR leading to an immune cold signalling environment compared to immune hot basal subtype of breast cancer.

We next sought to elucidate the mechanisms mediating this immune evasion. Tumors typically evade immune surveillance via two broad strategies: the systemic downregulation of antigen presentation (stealth) or the active suppression of infiltrating immune cells (exhaustion). Evaluation of key immune checkpoints and MHC pathways revealed a striking dichotomy between the TamR and basal states. The TamR state was fundamentally characterized by the dysregulation of the antigen presentation machinery (**Figure 6C, Supplementary Figure S5B**). We observed highly significant downregulation across the major histocompatibility complex (MHC) class I pathway, encompassing core MHC-I structural molecules, the peptide-loading complex, and broader antigen processing gene networks (**Figure 6C, Supplementary Figure S5B**). On the other hand, basal cell lines demonstrated classical features of active immune suppression, characterized by the significant upregulation of exhaustion and “don’t eat me” markers, including CD274 (PD-L1), NT5E, PVR, and CD47 (**Supplementary Figure S5C**). Conversely, these same markers were significantly downregulated in TamR cells compared to controls (**Supplementary Figure S5C**). Collectively, these data indicate that SERM-induced resistance does not rely on the expression of classical immune checkpoints to actively exhaust infiltrating T-cells; rather, it exhibits a “cold” microenvironment by reducing antigen presentation, potentially rendering the resistant cells functionally invisible to adaptive immune surveillance.

Given this dramatic shift in tumor-intrinsic immunogenicity, we next investigated the potential for broader paracrine remodeling of the tumor microenvironment (TME). To map these dynamics *in-silico*, we compared single cell transcriptomes of parental MCF7, Tamoxifen treated MCF7 and TamR cells (GSE195610) with a patient-derived scRNA-seq atlas (phs002371) (65) comprising non-malignant TME compartments, including stromal, endothelial, and diverse immune cell populations (**Supplementary Figure S5D**) (**see Methods**). By evaluating putative ligand-receptor (L-R) interactions between the tumor cells and the surrounding microenvironment, we identified distinct paracrine signaling interactions unique to the endocrine-sensitive and TamR states. We subsequently distilled these interactions into “sensitive” and “resistant” ligand signatures (**Supplementary Table 2**). Within the scRNA-seq trajectory, the expression of this resistant ligand signature progressively increased from the parental state through the intermediate persister phase, peaking in the fully resistant TamR cells (**Figure 6D**). The resistant ligand signature exhibited a robust positive correlation with the TamR pEMT score across all batch-corrected bulk datasets (**Figure 6E**), confirming that this co-expression pattern between the different ligand groups is inherently coupled to the hybrid lineage state. This paracrine reprogramming was driven by the selective upregulation of key immunomodulatory and pro-tumorigenic cytokines, most notably MIF and CXCL8, which are mediators of tumor invasiveness and metastatic niche formation (66,67). Finally, to determine the clinical potential of this therapy-induced microenvironmental remodeling, we evaluated our derived sensitive and resistant ligand signatures in patient cohorts. Using Kaplan-Meier survival analyses restricted strictly to luminal breast cancer subtypes, we observed a divergent prognostic impact: high expression of the “sensitive” ligand signature was significantly associated with a favorable prognosis (relapse free survival) and a lower hazard ratio (HR = 0.69, p-val < 0.001) (**Figure 6F**). In contrast, enrichment of the “resistant” ligand signature was highly predictive of poor clinical outcomes (relapse free survival), yielding a significantly elevated hazard ratio (HR = 1.28, p-val < 0.05) (**Figure 6F**). However, the “sensitive” ligand signature did not show significance in relapse free survival in basal (ER negative, HER2 negative) or HER2 positive (ER negative, HER2 positive) subtypes of breast cancer signifying the specificity of the “sensitive” signature towards luminal ER+ breast cancer patients (**Supplementary Figure S5E**). On the other hand, the “resistant” ligand signature exhibited a higher hazard ratio for relapse free survival in both basal and HER2+ breast cancer patients (**Supplementary Figure S5F**).

Taken together, these findings demonstrate that the pEMT–basal state stabilized by SERM resistance extends beyond tumor-intrinsic signaling, actively driving immune evasion via MHC-I silencing and coordinating a pro-tumorigenic, immunosuppressive and immune cold microenvironment that ultimately dictates adverse patient survival (**Figure 6G**).

**Supplementary Figure S5:**
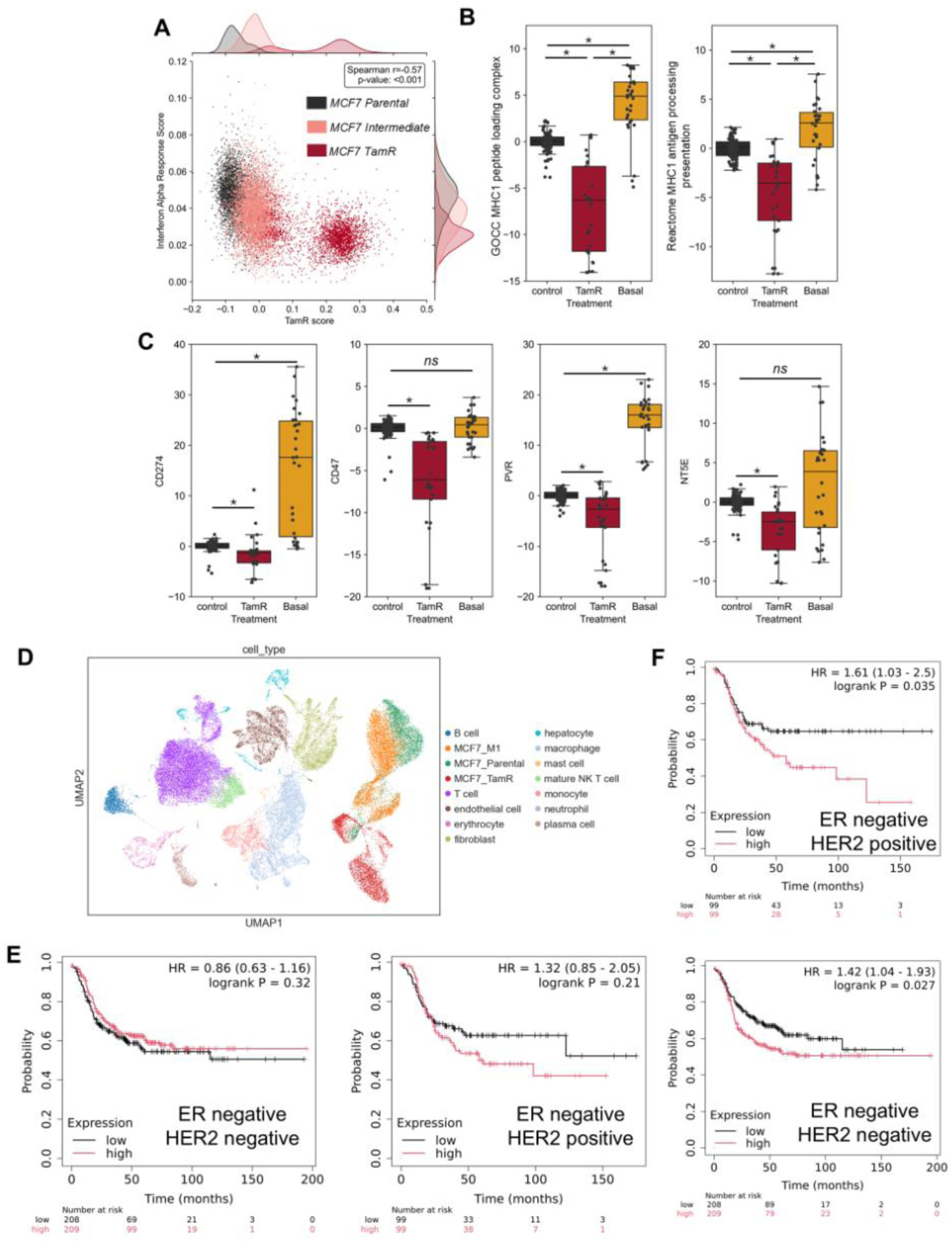
(A) Scatter plot showing single-cell level correlation between TamR-pEMT score and Hallmark interferon-α response score, with marginal density distributions and Spearman correlation indicated. (B) Boxplots showing ssGSEA scores for antigen presentation gene signatures: Gene Ontology MHC class I peptide loading complex (left) and MHC class I–mediated antigen processing and presentation (right) across control, TamR, and basal conditions. Statistical significance indicated by * (p < 0.05). (C) Boxplots showing normalized expression of CD274, CD47, PVR, and NT5E across control, TamR, and basal conditions. Statistical significance indicated by * (p < 0.05); ns denotes non-significant differences. (D) UMAP projection of integrated single-cell RNA-sequencing data showing cell lines (MCF7 parental, MCF7 intermediate and TamR) and non-malignant cell populations, with cell types annotated. (E) Kaplan–Meier survival analysis in Basal (left) and HER2+ (right) breast cancer patients stratified by “sensitive” ligand signature. (F) Kaplan–Meier survival analysis in HER2+ (top) and Basal (bottom) breast cancer patients stratified by “resistant” ligand signature.

## Discussion

Research into endocrine resistance in ER+ breast cancer has traditionally focused either on mechanisms that attenuate estrogen receptor (ERα) signaling and enable proliferative independence or on recurrent genomic alterations that sustain ligand-independent ER activity (68,69). Within this framework, resistance is commonly interpreted according to whether ER signaling is suppressed, restored, or bypassed (70,71), while comparatively less attention has been given to how the distinct mechanisms of action of endocrine therapies themselves shape adaptive cellular evolution. Endocrine agents do not simply reduce ER signaling to different extents; rather, they impose qualitatively distinct selective pressures through receptor antagonism, receptor degradation, or constitutive receptor activation. These mechanistic differences are likely to constrain and direct tumor adaptation along distinct transcriptional and epigenetic trajectories.

Using a batch-corrected integrative meta-analysis across multiple MCF7 endocrine treatment conditions, we systematically compared short-term endocrine perturbation states with long-term resistant states to define how sustained therapeutic pressure reshapes cellular identity. Acute endocrine exposure induced smaller magnitude transcriptional responses linked to altered ER signaling output, whereas long-term resistance was associated with more stable regulatory rewiring accompanied by widespread epigenetic remodeling, consistent with acquisition of adaptive drug-resistant states (72,73). Importantly, the trajectories of long-term adaptation differed between resistance modalities. Tamoxifen-resistant (TamR) cells exhibited the most extensive transcriptional and chromatin reprogramming, including coordinated luminal erosion, basal-like drift, enhancer redistribution, and acquisition of a pronounced partial epithelial–mesenchymal (pEMT) phenotype (**Figure 7A**). In contrast, fulvestrant-resistant (FulR) cells primarily showed suppression of ER-driven transcription with comparatively limited global reorganization, whereas ESR1 mutant models largely retained ligand-like ER hyperactivation programs (**Figure 7B**). Together, these findings demonstrate that endocrine resistance does not converge on a single resistant state but instead evolves along therapy-specific trajectories shaped by distinct selective constraints.

**Figure 7:**
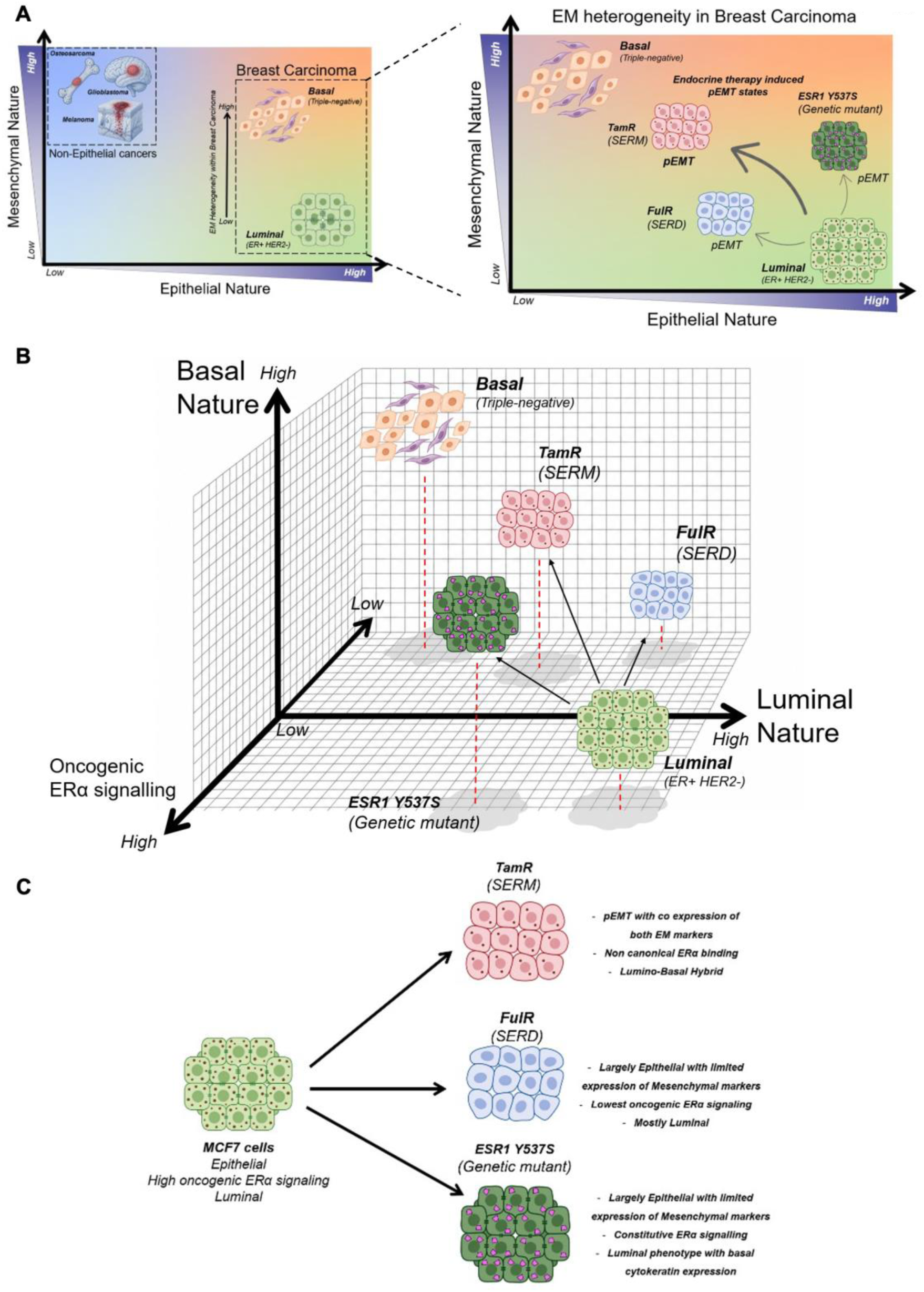
(A) Schematic showing EM heterogeneity in breast carcinoma. (Left) Two dimensional Epithelial-Mesenchymal plane showing the relative positions of breast carcinoma subtypes with respect to non-epithelial (mesenchymal origin) cancers such as glioblastoma, melanoma, osteosarcoma etc. (Right) Two dimensional Epithelial-Mesenchymal plane showing the stratification of tamoxifen- (TamR), fulvestrant (FulR)-resistant cancer and ESR1 Y537S mutant breast cancer as they get reprogrammed under selection pressure from different endocrine therapies. (B) Projection of TamR, FulR and ESR1 Y537S mutant cancer with the basal breast cancer on three-dimensional axes depicting luminal, basal and oncogenic ERα signalling. (C) Schematic showing EM, oncogenic signalling activity and lineage associated fates of TamR, FulR and ESR1 Y537S mutant breast cancer cells.

Importantly, ESR1-mutant resistance represents a phenotypically distinct adaptive state that should not be conflated with the lineage destabilization observed in TamR. ESR1 mutant models maintain hyperactive ER signaling while simultaneously expressing selective pEMT-associated programs and certain basal cytokeratins (25), as downstream consequences of persistent ER-driven transcriptional activity, including strong induction of canonical ER targets such as PGR (25). However, this phenotype differs fundamentally from the coordinated luminal erosion and epigenetic lineage reprogramming observed following prolonged SERM exposure (**Figure 7B**). Distinguishing these states required comparative meta-analysis across resistance modalities, as pathway enrichment or bulk transcriptional quantification in individual datasets alone would not adequately separate hyperactive ER-driven phenotypes from genuine lineage-transition states associated with destabilization of luminal regulatory architecture.

During mammary gland development, mammary stem cells and myoepithelial lineage that represent epithelial/epithelial-like populations that nonetheless frequently co-express epithelial and mesenchymal features, including simultaneous expression of markers such as CDH1 and SNAI2 (74). Similarly, basal breast cancers often occupy hybrid epithelial–mesenchymal states rather than fully mesenchymal states, retaining epithelial determinants while expressing basal lineage regulators such as TP63 together with varying degrees of EMT-associated transcriptional activity (53,74–76) (**Figure 7A**). Our findings suggest that endocrine therapy-induced plasticity in ER+ breast cancer partially converges toward this developmental and basal-like hybrid architecture. Under prolonged SERM exposure, luminal ER+ cells acquire selective basal and mesenchymal characteristics without undergoing complete lineage transdifferentiation (**Figure 7A**). This convergence toward a basal-pEMT state is most prominent in TamR models and may be linked to the altered activity of antagonized yet undegraded ERα retained during SERM treatment. By contrast, FulR cells lose oncogenic ER signaling but largely preserve core luminal lineage regulators such as GATA3 and FOXA1 (**Figure 7B**), indicating that suppression of ER activity alone is insufficient to induce extensive lineage destabilization. These observations support a more nuanced model of endocrine resistance in which adaptation is governed not only by changes in oncogenic signaling, but also by interactions between lineage-specifying transcriptional networks and epithelial–mesenchymal plasticity. Although these processes have often been studied independently (19,77), their coordinated contribution to endocrine-resistant state transitions had not previously been systematically resolved.

A major unresolved challenge in ER+ breast cancer biology is the limited efficacy of immunotherapy (78). ER+ tumors are typically immune cold, characterized by low immune infiltration and limited T-cell exhaustion, thereby reducing susceptibility to checkpoint blockade therapies (27,79,80). Our analysis provides a potential mechanistic explanation for why relapsed TamR tumors may remain immune cold despite acquisition of pEMT-associated characteristics. Classical EMT and pEMT programs are frequently associated with inflammatory signaling and enhanced immune infiltration (60), particularly in basal breast cancers (81), where increased immune engagement can subsequently generate exhausted T-cell populations targetable by immunotherapy (82) (**Figure 6G**). In TamR cells, however, acquisition of pEMT occurs concurrently with repression of IFNγ and IFNα signaling pathways, reduced antigen-presentation programs, and suppression of exhaustion-associated immune markers. Thus, although TamR cells acquire mesenchymal and basal-like transcriptional features, the accompanying epigenetic rewiring appears to uncouple pEMT from inflammatory immune activation. This suggests that therapy-induced pEMT in ER+ disease represents a biologically distinct state from the inflammatory EMT programs commonly observed in basal tumors. Maintenance or reinforcement of immune coldness through coordinated suppression of interferon-associated pathways may therefore contribute to the persistent resistance of endocrine-relapsed ER+ tumors to immunotherapeutic strategies.

Collectively, genetic and non-genetic resistance follow divergent evolutionary routes. ESR1 mutation reinforces ER signaling with partial lineage effects, FulR suppresses ERα without broad transcriptional reprogramming, and TamR coordinates epigenetic remodeling, luminal silencing, mesenchymal activation, and partial EM stabilization. These states can be conceptualized along three axes: oncogenic ER signaling intensity, lineage identity (luminal–basal), and epithelial–mesenchymal state. TamR shifts along all three axes, FulR primarily along ER signaling, and ESR1 mutants along ER signaling with limited EM remodeling (**Figure 7C**). This multi-axis framework provides a more nuanced model for endocrine-resistant heterogeneity.

An intriguing feature unique to TamR is activation of developmental and multi-tissue transcriptional programs, including but not limited to RUNX2- and SOX9-associated signaling pathways linked to osteogenic and chondrogenic differentiation (46,47). Engagement of skeletal developmental programs in TamR may therefore contribute to bone organotropism (83), suggesting that therapy-induced transcriptional alignment influences metastatic niche preference. Whether such lineage mimicry directly biases organ colonization remains an important open question.

Although our analysis integrates multiple datasets to systematically compare endocrine resistance modalities, comparable multi-omic datasets spanning short-term adaptation and long-term resistance remain limited across different ER+ model systems and therapeutic agents. Consequently, we were unable to comprehensively extend this framework across additional breast cancer cell lines or across multiple SERM and SERD compounds, such as raloxifene, lasofoxifene, elacestrant, or camizestrant. Future studies generating standardized transcriptomic and epigenomic datasets across diverse endocrine therapies, treatment durations, therapy dosing, and cellular backgrounds will be essential to determine which resistance mechanisms represent conserved modality-specific responses versus context-dependent adaptations. Such comparative analyses will be critical for defining how distinct drug mechanisms shape the evolutionary landscape of endocrine resistance.

In summary, SERM resistance represents an epigenetically stabilized partial epithelial–mesenchymal state intrinsically linked to partial basal lineage reprogramming and distinct from both SERD resistance and ESR1 mutational activation. By defining the basal–pEMT axis as a coordinated, chromatin-encoded module of therapy-induced plasticity, this work reframes endocrine resistance as a multidimensional evolutionary process rather than a single molecular endpoint.

## Methods

### Transcriptomic Data Acquisition and Pre-processing

Raw transcriptomic data, provided as Transcripts Per Million (TPM) counts aligned to the GRCh38.p13 reference genome, were retrieved for a specified set of Gene Expression Omnibus (GEO) datasets (listed in **Supplementary Table 1**). Gene identifiers were mapped to their corresponding HGNC gene symbols utilizing the GRCh38.p13 annotation file. Genes exhibiting zero expression across all samples within a dataset were excluded from downstream analyses. The resulting expression matrices were log-transformed using the formula log_2_(x+1).

### Cross-Study Harmonization and Batch Effect Correction

To facilitate integrated cross-study meta-analysis, the log-normalized expression matrices were parsed to identify and isolate the exact intersection of gene symbols present across all selected GSE datasets. A unified expression matrix consisting solely of these common genes was generated. To systematically correct for inter-study technical variations and batch effects, the harmonized matrices were processed using the COCONUT (COmbating CO-eNfounding Using conTrols) R package (84). The empirical Bayes-based correction was directed by the corresponding metadata “Condition” variable to appropriately partition control and treatment samples, without imposing platform-specific assumptions (byPlatform = FALSE).

### Sample Cohort Definition and Normalization

Post-batch correction, individual GSE datasets and their corresponding phenotypic metadata were concatenated into a singular, global data frame. The cohort was strictly filtered to include only samples originating from the MCF7 cell line or basal cell lines (BT20, BT549, CAL51, HCC1143, HCC1187, HCC1397, HCC38, HCC70, MDAMB157, and SUM149PT). Further stratification constrained the analysis to predefined experimental and treatment states: Control, Estradiol (E2), Tamoxifen (Tam), Tamoxifen-resistant (TamR), Fulvestrant (Ful), Fulvestrant-resistant (FulR), ESR1 Y537S mutant, and Basal phenotypes. To evaluate relative shifts in expression, the gene expression matrix was standardized via Z-score normalization computed relative to the baseline (control) cohort. Specifically, the mean and standard deviation of the control samples were calculated for each gene, and these baseline parameters were used to scale the entire common data frame.

### Single-Sample Gene Set Enrichment Analysis (ssGSEA)

Pathway enrichment was performed using the gseapy library (85) in Python to compute Single-Sample Gene Set Enrichment Analysis (ssGSEA) scores. Normalized Enrichment Scores (NES) were calculated across all samples using a predefined custom gene set library (**Supplementary Table 3**), with a minimum required gene set size of 5. The resulting ssGSEA NES matrix was subsequently Z-normalized relative to the control sample distribution, maintaining consistency with the gene-level expression normalization, to quantify relative pathway dysregulation.

### Non-Parametric Effect Size Evaluation and Statistical Testing

To robustly quantify the magnitude of expression shifts induced by different experimental conditions relative to the baseline (Control), Cliff’s Delta was computed for each gene. Cliff’s Delta was selected as a non-parametric measure of effect size, providing an evaluation of transcriptomic perturbations that is highly robust against outliers and does not assume normal distribution of the expression data. In parallel, the statistical significance of these differential expressions was assessed using the Mann-Whitney U test. For each condition, the mean and median of the Z-normalized expression values were also calculated. All resulting matrices (effect sizes, U-statistics, p-values, and central tendencies) were saved for downstream evaluations.

### Principal Component Analysis of Treatment Trajectories

To visualize global transcriptomic response trajectories across the different experimental conditions, Principal Component Analysis (PCA) was applied directly to the Cliff’s Delta effect size matrix. Utilizing effect sizes rather than raw or normalized expression values allows the PCA to capture axes of variation strictly driven by the magnitude of treatment response rather than baseline expression abundance. Prior to dimensionality reduction, any missing values within the effect size matrix were imputed to zero. A theoretical baseline ‘Control’ sample, defined by an effect size of zero across all genes, was appended to the matrix to securely anchor the origin of the multidimensional space. PCA was then computed using the first three principal components (n_components=3), and the multi-dimensional mapping of conditions was visualized using a customized 3D scatter plot.

### Feature Extraction and Gene Ontology Over-Representation Analysis

To isolate the molecular drivers shaping specific transcriptomic trajectories, the gene loadings for the principal components were extracted and sorted. For a targeted principal component, the top 2000 genes exhibiting the highest positive loadings and the 2000 genes with the strongest negative loadings were isolated as the primary contributors to that specific axis of variation (**Supplementary Table 3**). To elucidate the biological significance of these driving genes, Gene Ontology (GO) Over-Representation Analysis was performed. The extracted lists of top-contributing genes were queried against the PANTHER (Protein ANnotation THrough Evolutionary Relationship) (86) online classification system. Statistical enrichment of biological processes was determined using Fisher’s exact test. To stringently control for false positives across multiple comparisons, all resulting p-values were adjusted using the False Discovery Rate (FDR) correction method.

### Tracking Transcription Factor Regulatory Fates

To evaluate the transcriptomic fidelity and fate of specific transcription factor (TF) signatures across different phenotypic states, we systematically tracked the expression behavior of TFs between paired experimental conditions. First, the global effect size matrix was filtered to include only a predefined list of known human transcription factors. For a designated condition, TFs were stratified into up-regulated and down-regulated signatures relative to the baseline control, strictly filtering for genes that demonstrated a “large” or “medium” effect size magnitude based on Cliff’s Delta. The subsequent regulatory fate of these condition-specific TF signatures was then evaluated within a targeted “test” condition. Specifically, each TF originating from the source signature was queried in the test condition and re-classified into one of three distinct categories: structurally up-regulated (maintaining a large effect size with a positive Cliff’s Delta), down-regulated (large effect size with a negative Cliff’s Delta), or unchanged/small effect. The proportional distribution and absolute gene counts of these transcriptional fates were quantified and visualized using 100% stacked bar charts.

### Identification of pan-cancer epithelial and mesenchymal and breast cancer specific Estrogen signalling associated transcription factors and gene lists

To define robust, pan-cancer transcription factor (TF) signatures associated with epithelial-mesenchymal transition (EMT) states, gene-pathway correlation matrices derived from the Cancer Cell Line Encyclopedia (CCLE) were analyzed. Gene signatures for “Epithelial”, “Mesenchymal”, and “partial EMT” (pEMT) states were evaluated across multiple cancer types. TFs were retained for the core pan-cancer signatures if they exhibited a Pearson correlation coefficient(r) > 0.4 with the respective state signature in at least 6 distinct tissue types. At least 6 different tissue types were chosen so that we could capture genes that were specific to either epithelial or mesenchymal programs. For example, an epithelial TF is likely to be correlated to the epithelial program in only carcinoma as it is never/rarely expressed in purely mesenchymal tissue types. Having a higher number of tissues would penalise such TFs that are specific to and expressed only in either epithelial carcinomas or purely mesenchymal tissues. To identify breast-tissue-specific TFs strongly associated with estrogen signaling, CCLE gene-pathway correlation matrices for the Hallmark Estrogen signaling (Early and Late) pathways were calculated. For each TF, a tissue-specific Z-score was calculated to quantify its correlation within the breast lineage relative to the background distribution across all other available CCLE tissues. TFs were designated as breast-specific estrogen-responsive regulators if they demonstrated a breast lineage correlation(r) > 0.3 and a relative tissue Z-score(Z) > 1.5. A “Common ER” signature was also defined by taking the exact intersection of the filtered Early and Late estrogen response TF lists. Identical filtering criteria were used to also get full gene lists (not just transcription factors) as well.

### Tissue similarity Analysis

To contextualize treatment-induced transcriptional shifts within a broader cancer tissue type gene expression mimicry framework, global transcriptomic profiles from the Cancer Cell Line Encyclopedia (CCLE) were compared to the gene expression differences between control and COCONUT normalised gene expression across the different treat. Breast cancer cell lines were first annotated into Luminal and Basal subtypes. To quantify tissue-specific transcriptomic divergences against a biologically relevant baseline, Cliff’s Delta effect sizes were computed for pan-cancer tissue lineages relative to the Luminal breast cancer reference group within the CCLE group of cell lines. For statistical robustness, tissue cohorts were strictly filtered to retain those with at least 20 samples. This pan-cancer effect size matrix was subsequently merged with the COCONUT normalised condition-specific effect size matrix and strictly filtered to isolate known human transcription factors (TFs). This analysis is under the assumption that luminal cell lines in CCLE are transcriptionally similar to control MCF7 cell lines in our COCONUT normalised data as MCF7 is also a luminal cell line. Finally, to quantify global regulatory alignment, cosine similarity was calculated between the TF effect size vector of a selected treatment phenotype (e.g., TamR and FulR) and all other pan-cancer tissue gene expression profiles. This vector-based approach provided a standardized ranking of transcriptional mimicry, identifying which native tissue states most closely resemble the regulatory architecture driven by targeted therapeutic resistance.

### Single cell RNA seq analysis of Sensitive and Resistant MCF7 cells

Single-cell RNA sequencing data for MCF7 Parental, Tamoxifen-resistant (TamR), and an intermediate M1 state (GSE195610) were imported and concatenated using the scanpy Python framework. Following the removal of genes expressed in fewer than 3 cells, the raw count matrix was preserved, and the remaining data underwent total-count normalization (10,000 UMIs per cell) followed by log-transformation. Dimensionality reduction was performed by isolating highly variable genes, executing Principal Component Analysis (PCA), and constructing a neighborhood graph (10 neighbors, 40 principal components) prior to Uniform Manifold Approximation and Projection (UMAP) embedding. Functional pathway activity at single-cell resolution was evaluated utilizing two complementary approaches: the decoupler library was employed to compute Area Under the Curve (AUCell) enrichment scores directly on the raw counts using a custom breast cancer signature library (**Supplementary Table 3**). A composite phenotypic “TamR score” was subsequently derived for each cell by aggregating the positive signature modules and subtracting the negative modules, with the resulting continuous gradient scores mapped onto high-resolution UMAP visualizations.

### Estrogen Receptor Master Regulome Definition and Quantification of FOXA1 and GATA3 binding

To map the dynamic reorganization of the Estrogen Receptor (ER) cistrome during the acquisition of endocrine resistance, a “Master ER Universe” was established by merging all robustly identified ER binding sites (MACS2 narrowPeaks) from both MCF7 Parental and TamR conditions. This unified genomic coordinate map served as the baseline for all subsequent transcription factor co-occupancy analyses. To determine binary peak presence or absence within these defined ER boundaries, GATA3 and FOXA1 peak files from both cellular conditions were intersected with the Master ER Universe using bedtools, requiring a strict minimum overlap of 20% to call a locus “occupied.” Subsequently, to evaluate quantitative shifts in binding intensity at these sites, read counts were extracted from respective smoothed CPM BigWig files across all three factors (ER, FOXA1, GATA3) in both conditions. Binding dynamics were mathematically categorized by computing the log_2_ fold change (L2FC) of signal intensity (TamR vs. MCF7), with a stringent threshold of 0.58 (equivalent to a 1.5-fold change) used to classify sites as significantly gained, lost, or structurally stable. Finally, these mapped and quantitatively scored regulatory regions were annotated to their nearest protein-coding genes via genomic proximity utilizing the GRCh38 reference annotation. This allowed for the aggregation of localized enhancer and promoter binding dynamics (e.g., mean L2FC across all ER/FOXA1/GATA3 sites mapped to a specific gene) to track the targeted regulatory trajectory of predefined phenotypic gene sets (Epithelial, Mesenchymal, and ER-associated TFs) during the resistant transition.

### Single-Cell Integration and Intercellular Communication Analysis

To investigate intercellular communication between distinct breast cancer phenotypic states and the tumor microenvironment (TME), *in-vitro* single-cell RNA-sequencing profiles of MCF7 Parental, M1, and Tamoxifen-resistant (TamR) cells (GSE195610) were integrated with a primary ER+/HER2- patient cohort (phs002371). Patient data were strictly filtered to isolate non-malignant TME populations and restricted to protein-coding genes. Following dataset concatenation across common intersecting genes, the unified matrix was preprocessed by removing genes detected in fewer than 10 cells, normalizing to 10,000 counts per cell, and applying a log-transformation. Dimensionality reduction was achieved through data scaling, Principal Component Analysis (PCA) on highly variable genes, and UMAP embedding based on a 15-neighbor graph utilizing 30 principal components. Ligand-receptor crosstalk was subsequently quantified using the LIANA framework (87) coupled with the OmniPath consensus resource, applying a minimum cellular expression threshold of 10%. Directional communication networks were modeled from the specific MCF7 source states to distinct immune and stromal TME target populations. Putative paracrine interactions were filtered for statistical stringency utilizing a specificity rank threshold of ≤ 0.05, allowing for the extraction and comparative evaluation of condition-specific signaling axes, specifically identifying unique receptor-ligand pairs emergent in the TamR state relative to the Parental baseline.

### HiC data processing for polymer modeling

We constructed the three-dimensional organization of a 12 Mb chromatin region containing the GATA3 gene using a bead-spring polymer model. The modeled region spans genomic coordinates 3 Mb to 15 Mb on chromosome 10 and was analyzed for the MCF7, TamR, and FulR cell lines at 10 kb resolution. The corresponding HiC datasets were obtained from GSE118712 (7) for all three cell lines at the resolution of 10 kb. To identify the chromosomal compartments, the available PC1 values were used. Homer ndHiCCompartments.pl script was used to identify the continuous regions of positive and negative PC1 values (88). Positive PC1 values corresponded to compartment A (active/euchromatin region) and negative values represented compartment B (inactive/heterochromatin region).

### Polymer simulations

Molecular dynamics simulations were performed to generate three-dimensional structural ensembles of the 12 Mb chromatin region for the MCF7, TamR, and FulR cell lines containing the GATA3 gene. The chromatin segment was modeled using the Kremer-Grest bead-spring polymer model, where the polymer chain consists of 1200 beads, each representing 10 kb of genomic DNA. In this model, the GATA3 genomic region was defined such that its transcription start site (TSS) is explicitly mapped onto the polymer, corresponding to the 506th bead in a 1200-bead chain. Super-enhancer (SE) regions were identified within a genomic window spanning 3–15 Mb surrounding the locus.

Polymer beads interact via a detailed particle-interaction potential commonly used in polymer simulations, consisting of three components:

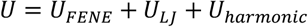

The bonded beads interact with finite extensible nonlinear elastic (FENE) potential (89)

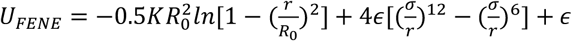

with length constant equal (𝑅_0_) to 1.6𝜎 and strength 𝐾 of 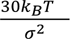. To account for the excluded volume potential between any two non-bonded beads, the repulsive part of the lennard-jones (LJ) potential

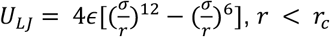

is used with 𝜎 is the length scale unit, 𝜖 = 𝑘_𝐵_𝑇 is the energy scale unit, and 𝑟_𝑐_ is the interaction range cutoff (1.12𝜎). The non-bonded beads are connected with harmonic constraint 𝑈_𝐻𝑎𝑟𝑚𝑜𝑛𝑖𝑐_ = 𝑘(𝑟 − 𝑟_0_)^2^, where the value of k depends on the contact probability c_ij_ between i^th^ and j^th^ bead, and k is defined as k=k c ^2^ with k_0_ = 0.05 (90,91).

The bead-spring polymer model was simulated using the LAMMPS simulation package (92). The equations of motion were integrated using the Verlet algorithm, while the temperature was maintained at a constant value using a Langevin thermostat set to T=1.0 in reduced units. The friction coefficient γ, related to the solvent viscosity η through γ=6πηR, where R is the particle radius, was set to 1.0.

Simulations were initialized from a self-avoiding random walk configuration, and the system was equilibrated in the NVT ensemble for 10^7^ timesteps. After equilibration, trajectories were saved at regular intervals, yielding 2×10^4^ distinct configurations for the chromatin region in each simulation. To improve sampling, simulations were repeated with 20 independent initial configurations, resulting in a total of 4×10^5^ configurations. These configurations were used to compute the contact maps and perform further analyses. To evaluate the accuracy of the model, we calculated the Pearson correlation coefficient between the simulated contact maps and the corresponding experimental Hi-C contact maps (93–95). The resulting correlation values ranged from 0.72 for the MCF7, 0.7 for TamR, and FulR cell lines, indicating good agreement between the simulated and experimental chromatin contact patterns and validating the polymer modeling approach.

### Analysis of simulation of trajectories

All chromatin configurations obtained from the polymer simulation trajectories were analyzed to quantify the compartment segregation and kinetics of super-enhancer and promoter (SE-P) interactions.

Compartmentalization was quantified using a segregation ratio defined as:

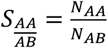

where 𝑁_𝐴𝐴_ and 𝑁_𝐴𝐵_ denote the average number of A-A and A-B contacts, respectively. Here, 𝑁_𝐴𝐴_ quantifies interactions between loci belonging to the same compartment (A), while 𝑁_𝐴𝐵_ captures interactions between loci from different compartments (A and B). A higher value of 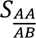 indicates stronger compartmental segregation. We also quantified the cluster size around the promoter by considering both super-enhancers and non-super-enhancers within a cutoff distance of 3σ, in order to assess how compartment switching influences the specificity of promoter interactions.

To characterize the SE-P binding kinetics, we calculated the binding and unbinding rates for each SE-P pair using all 20 independent simulation trajectories. The binding rate (𝑏_𝑖𝑗_) was defined as the inverse of the mean time required for a super-enhancer to associate with the promoter within a cutoff distance of 3σ. Conversely, the unbinding rate (𝑢_𝑖𝑗_) was defined as the inverse of the mean dwell time for which the super-enhancer remains associated with the promoter before dissociation (96)

For each SE–P pair, the dwell time distributions were fitted using a double-exponential function of the form

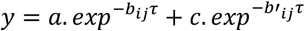

Since the slower process is expected to be rate-limiting, the corresponding slow decay constants obtained from the fits were used to estimate the effective binding (𝑏_𝑖𝑗_) and unbinding (𝑢_𝑖𝑗_) rates for the interaction between super-enhancer i and promoter j (91). The residence time (𝜏) for each SE–P pair was calculated as the time the promoter remains in the bound state, given by

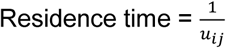

Based on residence time, super-enhancers were ranked from most to least effective in order to assess the impact of compartment organization on their specific interactions with the promoter.

## Author contributions

Conceptualized and designed research: MKJ, SS; supervised research: MKJ, SH, HK; performed research: SS, SK, SSe; funding acquisition: MKJ, SH, HK; manuscript writing/editing: SS (prepared first draft). All authors contributed to data analysis, review and editing of the manuscript.

## Acknowledgments

S.S. is supported by PMRF (Prime Ministers’ Research Fellowship) awarded by Government of India. M.K.J. is also supported by Param Hansa Philanthropies. H.K. acknowledges the financial support from the Department of Science and Technology, India, under “Fund for Improvement of S&T Infrastructure (SR/FST/PS-I/2020/140)” scheme. This research was funded, in part, by U.S. National Cancer Institute grant 1-ZIA-BC012176-02 to S.H. This research was supported [in part] by the Intramural Research Program of the National Institutes of Health (NIH). The contributions of the NIH author(s) are considered Works of the United States Government. The findings and conclusions presented in this paper are those of the author(s) and do not necessarily reflect the views of the NIH or the U.S. Department of Health and Human Services.

During the preparation of this work, the authors used Google Gemini and ChatGPT in order to improve the readability of the manuscript. After using this tool, the authors reviewed and edited the content as needed and take full responsibility for the final content of the publication.

